# Mitotic checkpoint gene expression is tuned by coding sequences

**DOI:** 10.1101/2021.04.30.442180

**Authors:** Eric Esposito, Douglas E. Weidemann, Jessie M. Rogers, Claire M. Morton, Erod Keaton Baybay, Jing Chen, Silke Hauf

**Affiliations:** Department of Biological Sciences and Fralin Life Sciences Institute, Virginia Tech, Blacksburg, VA 24061, USA

**Keywords:** mitosis, spindle assembly checkpoint, mRNA decay, gene expression noise

## Abstract

The mitotic checkpoint (also called spindle assembly checkpoint, SAC) is a signaling pathway that safeguards proper chromosome segregation. Proper functioning of the SAC depends on adequate protein concentrations and appropriate stoichiometries between SAC proteins. Yet very little is known about SAC gene expression. Here, we show in fission yeast (*S. pombe*) that a combination of short mRNA half-lives and long protein half-lives supports stable SAC protein levels. For the SAC genes *mad2^+^* and *mad3^+^*, their short mRNA half-lives depend on a high frequency of non-optimal codons and mRNA destabilization mediated through the RNA helicase Ste13 (*S.c.* Dhh1). In contrast, *mad1^+^* mRNA half-life is short despite a relatively high frequency of optimal codons and despite the lack of known destabilizing motifs in its mRNA. Hence, although they are functionally related, different SAC genes employ different strategies of expression. Taken together, we propose that the codon usage of SAC genes is fine-tuned for proper SAC function. Our work shines light on the gene expression features that promote spindle assembly checkpoint function and suggests that synonymous mutations may weaken the checkpoint.

## Introduction

The spindle assembly checkpoint (SAC; also called mitotic checkpoint) is a eukaryotic signalling pathway that delays cell cycle progression when chromosomes have not yet become properly attached to microtubules during mitosis (Kops *et al*, 2020; Lara-Gonzalez *et al*, 2012; Musacchio, 2015). Proper function of the SAC needs adequate SAC protein concentrations (both too low or too high expression can be detrimental) and needs adequate stoichiometries between proteins in the pathway (Chung & Chen, 2002; Gross *et al*, 2018; Heinrich *et al*, 2013; Ryan *et al*, 2012; Schuyler *et al*, 2012). This makes it important to quantitatively understand SAC gene expression. Yet, expression of these genes has not been studied in any detail.

The protein network of the SAC, on the other hand, is well understood. While the SAC is active, it forms the mitotic checkpoint complex (MCC), which prevents the anaphase-promoting complex (APC/C) from initiating anaphase (Pines, 2011). A key effector of the SAC is the Mad1/Mad2 complex, a tetramer of two Mad1 and two Mad2 molecules (Chen *et al*, 1999; Sironi *et al*, 2002). Mad1 homodimerizes through a long, parallel inter-molecular coiled-coil at its N-terminus, which is followed by the Mad2-binding motif and a C-terminal RWD domain (Chen *et al.*, 1999; Kim *et al*, 2012; Piano *et al*, 2021; Sironi *et al.*, 2002). The Mad1-binding partner Mad2 is a HORMA domain protein (named after Hop1, Rev7 and Mad2) that can change its conformation between open (O) and closed (C) (Aravind & Koonin, 1998; Luo *et al*, 2002; Luo *et al*, 2004). When Mad2 binds Mad1, the C-terminus of Mad2 wraps around the Mad1 polypeptide similar to a seat belt and Mad2 adopts the closed conformation (Fig. 1E) (Luo *et al.*, 2002; Sironi *et al.*, 2002). This results in a tight complex with no measurable off-rate *in vitro* (Chen *et al.*, 1999; Sironi *et al*, 2001; Vink *et al*, 2006). If and to what extent formation of the intricate Mad1/Mad2 complex is aided by other factors is unknown.

**Figure 1.**
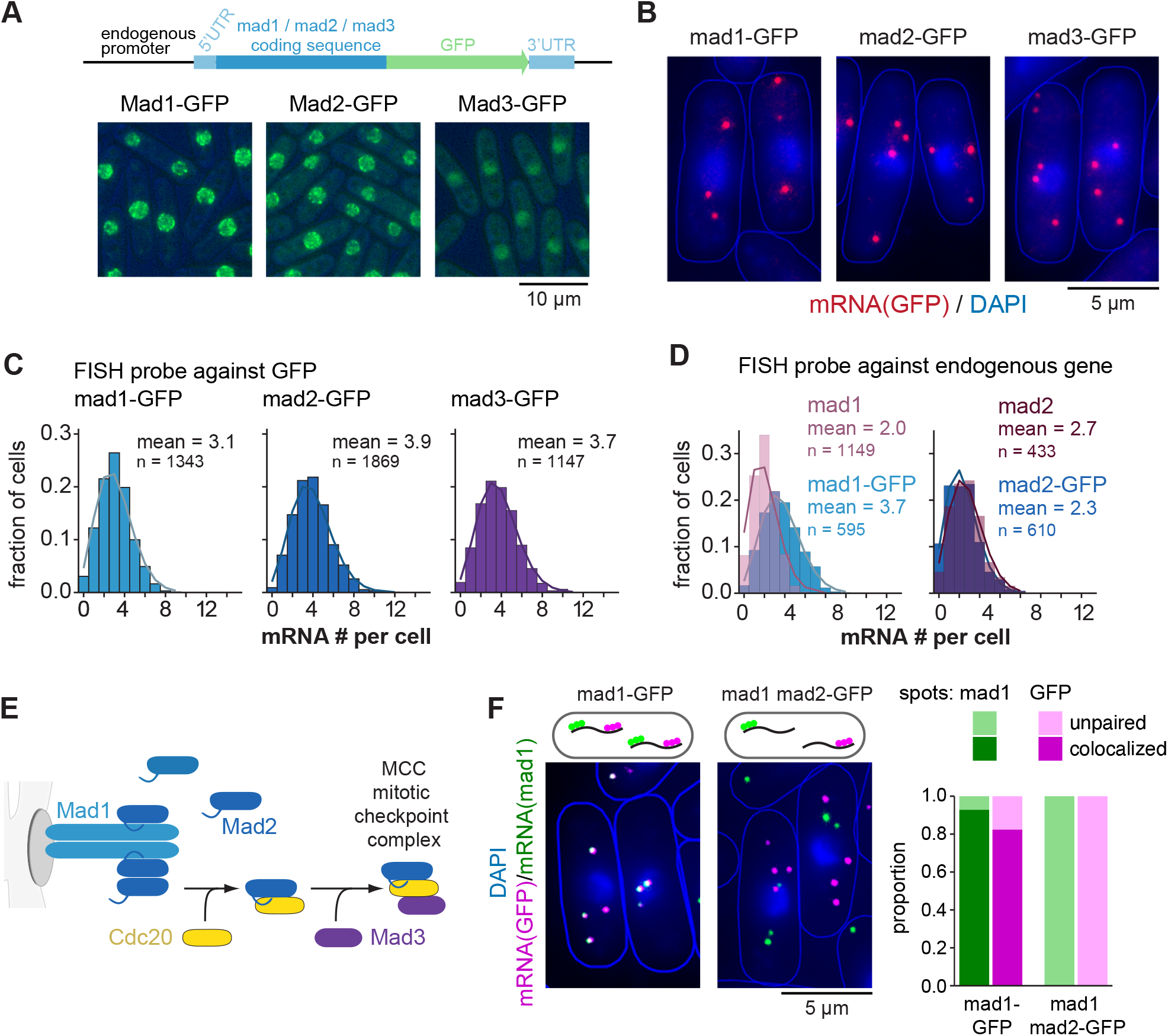
Low steady-state mRNA numbers of checkpoint genes *mad1*, *mad2* and *mad3*. **(A)** Schematic of marker-less GFP-tagging at the endogenous locus and representative live cell images of Mad1-, Mad2-, and Mad3-GFP strains (average intensity projections). **(B)** Representative images of single molecule mRNA FISH (smFISH) staining using probes against GFP (red). DNA was stained with DAPI (blue). The gamma-value was adjusted to make the cytoplasm visible; cell shapes are outlined in blue. **(C)** Frequency distribution of mRNA numbers per cell determined by smFISH. Curves show fit to a Poisson distribution. Two replicate experiments were combined (shown individually in Figure S1). **(D)** Frequency distribution of mRNA numbers per cell using FISH probes against the endogenous genes and using either strains expressing the GFP-tagged gene or the endogenous, untagged gene. **(E)** Overview of the interactions between Mad1, Mad2 and Mad3. **(F)** Co-staining by smFISH using probes against mad1 and GFP either in a strain expressing *mad1-GFP* or in a strain expressing wild-type *mad1* and *mad2-GFP*. Cytoplasmic mad1 (green) or GFP RNA spots (magenta) were quantified as co-localizing or not with the respective other. For the *mad1-GFP* strain, 565 cells that had both mad1 and GFP spots and a total of 1,637 mad1 spots and 1,848 GFP spots were analyzed; 30 cells were not considered since they either only contained mad1 or only GFP spot(s). For the *mad1 mad2-GFP* strain, 658 cells that had both GFP and mad1 spots and a total of 1,165 mad1 spots and 1,643 GFP spots were analyzed; 73 cells were not considered since they either only contained GFP or only mad1 spot(s).

Through a different surface, Mad2 can form heterodimers between its two conformations (O-C) (Mapelli *et al*, 2007). Dimerization of Mad1/C-Mad2 with O-Mad2 facilitates binding of this O-Mad2 molecule to the APC/C activator Cdc20 (Slp1 in *S. pombe*). O-Mad2 changes its conformation in the process, forming C-Mad2/Cdc20 through the same seat belt type of binding (Luo *et al.*, 2002). Subsequent binding of BubR1 (Mad3 in yeast) to C-Mad2/Cdc20 results in the mitotic checkpoint complex (MCC) (Chao *et al*, 2012; Sudakin *et al*, 2001). The MCC then inhibits the APC/C to block anaphase (Alfieri *et al*, 2016; Pines, 2011).

Because the SAC plays a central role in preventing chromosome mis-segregation and because persistent chromosome mis-segregation is a driver of tumor evolution, SAC malfunction is suspected to contribute to carcinogenesis (Funk *et al*, 2016; Gordon *et al*, 2012). Mouse models have shown that impairing the SAC promotes chromosome mis-segregation and tumor formation (Baker *et al*, 2005; Holland & Cleveland, 2009; Schvartzman *et al*, 2010). Completely abolishing the SAC, however, is detrimental to human cells (Dobles *et al*, 2000; Kops *et al*, 2004; Michel *et al*, 2004; Schukken *et al*, 2021), and suppression of the SAC may in fact be a successful therapeutic strategy against some cancer types (Cohen-Sharir *et al*, 2021; Quinton *et al*, 2021). Together, these results indicate that tuning SAC function can make the difference between normal growth, cancerous growth and cell death.

Although the SAC network has been studied in much detail from a protein-centric view, little is known about SAC gene expression. Understanding this regulatory layer is important, because changes in SAC protein concentrations can cause SAC malfunction—at least partly because proper stoichiometries, such as between Mad1 and Mad2, are important for function (Chung & Chen, 2002; Gross *et al.*, 2018; Heinrich *et al.*, 2013; Ryan *et al.*, 2012; Schuyler *et al.*, 2012). Here, using fission yeast (*S. pombe*), we study the mRNA layer of SAC gene expression and provide evidence that a combination of short mRNA and long protein half-lives ensures a stable concentration of SAC proteins over time and between cells. How mRNA half-life is controlled is still poorly understood. Recently, codon optimality of the mRNA has been implicated in influencing half-life (Boël *et al*, 2016; Presnyak *et al*, 2015). Our findings indicate that codon usage bias in *mad2^+^* and *mad3^+^*, but not *mad1^+^*, contributes to their short mRNA half-lives, and that the coding sequence of *mad1^+^* carries alternative features that influence expression of this gene. Overall, our findings shine light on gene expression features that promote SAC function and raise the possibility that synonymous mutations may impair the SAC.

## Results

### SAC mRNA numbers are Poisson distributed around means of two to four per cell

We previously quantified the concentration of SAC proteins fused to GFP in *S. pombe* and determined protein concentrations in a range between 30 and 150 nM with strikingly little inter-cell variability (i.e. low ‘noise’) (Heinrich *et al.*, 2013). In these strains, GFP had been fused by traditional tagging, changing the endogenous 3’ UTR to that of the *S. cerevisiae ADH1* gene and appending an antibiotic-resistance gene, which both may alter gene expression. To avoid such effects, we now employed CRISPR/Cas9-mediated scarless genome editing (Jacobs *et al*, 2014). We fused ymEGFP (yeast codon-optimized, monomeric enhanced GFP; in the following just ‘GFP’) to the SAC genes *mad1^+^*, *mad2^+^* and *mad3^+^* without any change to the surrounding sequences (Fig. 1A). Immunoblots showed concentrations broadly similar to the previous strains (Fig. S1A) and strains were not sensitive to the microtubule drug benomyl, suggesting that SAC functionality was maintained (Fig. S1B).

The mean SAC mRNA numbers per cell, determined by single-molecule mRNA FISH with probes targeting GFP, were in the range of 3 to 4, even lower than the means of 4.5 to 6 that we had previously observed (Fig. 1B,C; S1C) (Heinrich *et al.*, 2013). This indicates that the traditional tagging strategy indeed influenced gene expression. To test whether expression in the new strains resembles endogenous expression, we used FISH probes against endogenous *mad1^+^* and *mad2^+^* and compared strains expressing the endogenous untagged gene with strains expressing the GFP-tagged gene. For *mad2^+^*, the efficiency of the gene-specific probe was lower than the GFP probe (Fig. S1D), but the mean mRNA number for untagged and tagged *mad2^+^* was comparable (Fig. 1D). For *mad1^+^* in contrast, the *mad1^+^* probe closely resembled the GFP probe in efficiency (Fig. S1D), but untagged *mad1^+^* showed even fewer mRNA molecules than *mad1-GFP* (Fig. 1D). This suggests that the mere addition of GFP, without any changes in the UTRs or surrounding sequences can change expression of *mad1^+^*.

While the mean mRNA numbers per cell for the GFP tagged genes were in the range of 3 to 4, the numbers in single cells ranged vastly, from 0 to around 9 (Fig. 1C,D). As expected (Padovan-Merhar *et al*, 2015; Sun *et al*, 2020; Zhurinsky *et al*, 2010), smaller cells had on average lower numbers than larger cells (Fig. S1E). However, even cells of the same size could differ in mRNA number by 8 or more (Fig. S1E). The spread of mRNA numbers in the cell population was well approximated by a Poisson distribution (Fig. 1C,D). A Poisson distribution is expected from constitutive expression, where mRNA is synthesized and degraded in uncorrelated events but with a uniform probability over time. In contrast, “bursty” expression (characterized by alterations of promoter activity and inactivity) would result in an even wider distribution (Zenklusen *et al*, 2008). These results therefore indicate that SAC mRNA numbers vary considerably, but that this variation is within the expected range for constitutive expression.

### *mad1^+^* and *mad2^+^* mRNAs do not co-localize in the cytoplasm

The mRNA FISH data also provides the location of mRNAs. Recent work has suggested that co-translational assembly of protein complexes is more prevalent than previously thought (Schwarz & Beck, 2019). How the stable Mad1/Mad2 complex assembles is unknown. When heterodimeric complexes assemble while both subunits are being translated, their mRNAs will co-localize (Panasenko *et al*, 2019). We asked whether this is the case for Mad1 and Mad2. We stained endogenous *mad1^+^* mRNA (using a *mad1^+^* probe) and *mad2^+^-GFP* mRNA (using a *GFP* probe). While the *mad1^+^* and *GFP* probes showed strong co-localization in a *mad1-GFP* strain, there was no evidence for co-localization of *mad1^+^* and *mad2^+^-GFP* mRNA (Fig. 1F). This absence of mRNA co-localization excludes that the Mad1/Mad2 complex forms by synchronous co-translational assembly. We will discuss other possibilities below.

### Low protein noise can be explained through long protein and short mRNA half-lives

To analyze if and to what extent the strong mRNA variation propagates to the protein level, we quantified GFP-tagged Mad1, Mad2 and Mad3 in single cells using our ‘Pomegranate’ image analysis pipeline, which allows for 3D segmentation (Fig. S2, S3A) (Baybay *et al*, 2020). To subtract autofluorescence, we mixed the GFP-expressing cells with cells not expressing GFP (Fig. S2). Unlike for the mRNA, we observed little cell-to-cell variability in the SAC protein concentrations (Fig. 2A). As a comparison we imaged a ‘noisy’ *S. pombe* protein, Nmt1 (Saint *et al*, 2019), which indeed showed pronounced cell-to-cell variability (Fig. 2A, S2C). A measure of variability is the coefficient of variation (CV; standard deviation divided by mean). The CV for Mad1-, Mad2- or Mad3-GFP was in the range of 0.15 to 0.2, whereas that for Nmt1-GFP was around 0.5 to 0.75 (Fig. 2A).

**Figure 2.**
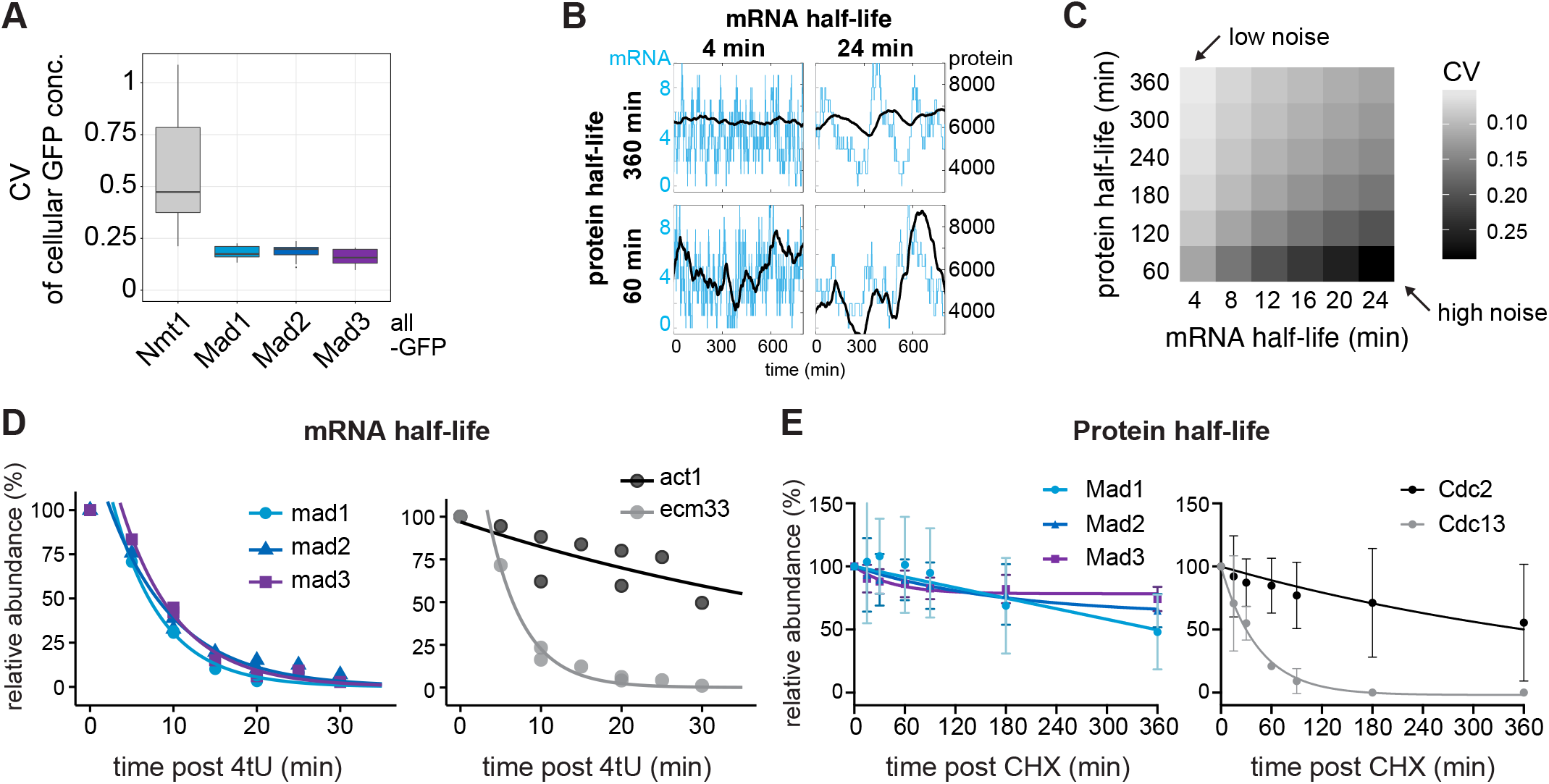
The checkpoint genes *mad1*, *mad2* and *mad3* combine short mRNA and long protein half-lives, explaining low noise. **(A)** Cellular noise (coefficient of variation, CV = std / mean) in live-cell microscopy images (Nmt1-GFP: n = 7; Mad1-GFP: n = 11; Mad2-GFP: n = 20; Mad3-GFP: n = 10 images; single images had between 16 and 79 GFP-positive and between 7 and 94 GFP-negative (control) cells). **(B)** Stochastic gene expression simulations (see Methods) from selected mRNA/protein half-life combinations (corners of the square shown in C). Synthesis rates were set to obtain a mean mRNA number of 4 per cell, and a mean protein number of 6000 per cell. **(C)** Predicted coefficient of variation (CV = std / mean) of the protein number per cell, assuming different mRNA and protein half-lives, using the same underlying model as in B. Synthesis rates were adjusted to maintain a mean mRNA number per cell of 3.5, and a mean protein number per cell of 6000 (approx. 100 nM). **(D)** mRNA abundances by qPCR following metabolic labeling and removal of the labeled pool (two independent experiments). Lines indicate fit to one-phase exponential decay, excluding the measurements at t = 0 in order to accommodate for non-instantaneous labeling by 4tU. *Act1* and *ecm33* were used as long and short half-life controls, respectively; qPCR for the endogenous mRNAs. **(E)** Protein abundances after translation shut-off with cycloheximide (n = 3 experiments, error bars = std). Lines indicate fit to a one-phase exponential decay. Cdc2 and Cdc13 were used as long and short half-life controls, respectively. Immunoblots for the endogenous proteins (no tag). One representative experiment shown in Figure S3.

This raised the question how the protein concentrations of Mad1, Mad2 and Mad3 can be homogeneous across the population when the mRNA numbers are highly variable. To identify the underlying mechanism, we considered different rates of transcription and translation, as well as of mRNA and protein degradation, that all yield mRNA and protein numbers similar to those that we observe for *mad1^+^*, *mad2^+^* and *mad3^+^* (see Methods for details). The longer the mRNA half-life, the longer a state of low or high mRNA numbers persists; and the shorter the protein half-life, the more closely protein concentrations follow the mRNA numbers (Fig. 2B). Hence long mRNA half-lives and short protein half-lives favour noise, whereas short mRNA-half lives and long protein-half-lives suppress noise (Fig. 2B,C; S3B). In the latter case, the long persistence time of proteins averages out fast fluctuations at the mRNA level (Fig. 2B).

To ascertain whether this prediction is met by SAC genes, we measured mRNA and protein half-lives. We determined mRNA half-life by metabolic labelling followed by depletion of the labelled pool and quantification of the remaining pool by quantitative PCR. The mRNA half-lives for *mad1^+^*, *mad2^+^* and *mad3^+^* were all in the range of a few minutes (*mad1^+^*: 3.8 min, *mad2^+^*: 6.4 min, *mad3^+^*: 4.1 min) (Fig. 2D; S5E, S6E). This was consistent with half-lives of 3 to 5 min for these genes in a large-scale study using metabolic labelling (Fig. S3D) (Eser *et al*, 2016). RNA half-lives have been notoriously difficult to measure, with much variability between studies (Carneiro *et al*, 2019). An earlier *S. pombe* study (Hasan *et al*, 2014) found longer half-lives across the entire transcriptome, but even in this study SAC genes were at the lower end of mRNA half-lives (Fig. S3D). As controls, we measured two unrelated genes with reportedly long and short half-life (Eser *et al.*, 2016), *act1^+^* and *ecm33^+^*, which behaved as expected (Fig. 2D). We determined protein half-lives by translation shut-off using cycloheximide, followed by immunoblotting. The half-lives of Mad1, Mad2 and Mad3 were in the range of many hours, considerably longer than the typical *S. pombe* cell cycle of 2.5 hours (Fig. 2E; S3E) and broadly consistent with previous data (Christiano *et al*, 2014; Horikoshi *et al*, 2013; Sczaniecka *et al*, 2008). This large difference in mRNA and protein half-lives explains the low cell-to-cell variability in protein concentration despite the considerable variation in mRNA numbers (Fig. 2C). The short mRNA half-life is therefore important to mitigate the effect of the large variation in mRNA numbers.

### *mad2^+^* and *mad3^+^* have low codon stabilization coefficients

One of the determining factors for mRNA half-life is codon optimality, which positively correlates with mRNA stability in several eukaryotes (Hanson & Coller, 2018). The codon stabilization coefficient (CSC) describes the correlation between the occurrence of a codon in mRNA transcripts and experimentally determined mRNA stability (Presnyak *et al.*, 2015). The CSC for a codon is positive if this codon is over-represented in stable mRNAs and negative if over-represented in unstable mRNAs. Similar to Harigaya and Parker (Harigaya & Parker, 2016), we determined CSC values for *S. pombe* based on large-scale mRNA half-life measurements (Eser *et al.*, 2016; Hasan *et al.*, 2014). The CSC value for each gene (CSC_g_) is the arithmetic mean of the CSC values of all codons in that gene. As had been seen before (Harigaya & Parker, 2016; Presnyak *et al.*, 2015), the CSC_g_ correlated with other measures of codon optimality such as the percentage of optimal codons or the tRNA adaptation index (tAI) (Fig. S4A). Since the SAC genes had short mRNA half-lives, we expected them to have low CSC_g_ values. Indeed, *mad2^+^* and *mad3^+^* were among the 20% of protein-coding genes with the lowest CSC_g_ values (Fig. 3A,B). This result was independent of which large-scale mRNA half-life data or which correlation parameter was used (Fig. S4C,D). These results raise the interesting possibility that codon usage in *mad2^+^* and *mad3^+^* contributes to their short mRNA half-life. The *mad1^+^* gene showed different characteristics, which we will discuss below.

**Figure 3.**
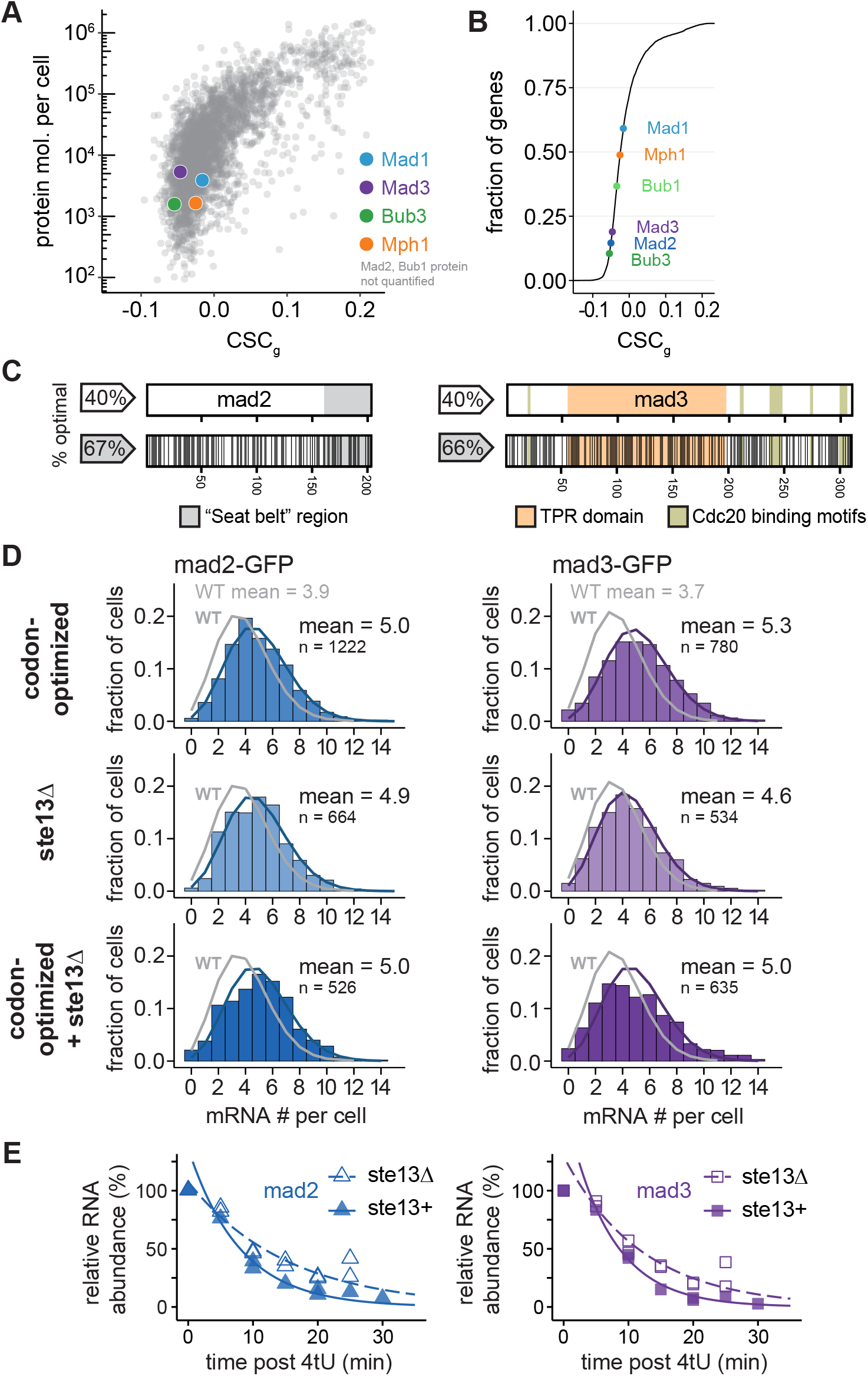
Codon optimization and *ste13* deletion increase the steady-state mRNA numbers of *mad2* and *mad3*. **(A)** The mean CSC value for each *S. pombe* gene (CSC_g_) relative to protein number per cell determined by mass spectrometry (Carpy *et al.*, 2014). CSC for each codon was determined using the mRNA half-life data by Eser et al. (Eser *et al.*, 2016) as described in Methods. Colored dots highlight proteins of interest. For Mad2 and Bub1, no protein abundance data was available. **(B)** Cumulative frequency distribution of the CSC_g_ values for protein-coding *S. pombe* genes. The position of spindle assembly checkpoint genes is highlighted. **(C)** Schematic of the *mad2* and *mad3* genes. Regions coding for important structural features are highlighted. Black lines in the bottom graph indicate synonymous codon changes in the codon-optimized version. **(D)** Frequency distribution of mRNA numbers per cell determined by mRNA FISH. Probes were against the GFP portion of the respective fusion gene. Curves show fit to a Poisson distribution. Grey curves show the fit from wild-type *mad2-GFP* or *mad3-GFP* cells (WT, Figure 1). Two replicate experiments were combined for codon-optimized genes and *ste13* deletion (shown individually in Figure S3). **(E)** Time course of mRNA abundances by qPCR following metabolic labeling and removal of the labeled pool (two independent experiments). Lines indicate fit to one-phase exponential decay, excluding measurements at t = 0 in order to accommodate for non-instantaneous labeling by 4tU. The *ste13^+^* data are the same as in Figure 2.

### Codon-optimization and *ste13^+^* deletion increase mRNA concentration and mRNA half-life of *mad2^+^* and *mad3^+^*

To test if codon usage contributes to the short mRNA half-lives, we codon-optimized *mad2* and *mad3* and inserted the codon-optimized sequence at the respective endogenous locus (Fig. 3C; S4F). The GFP tag, which remained unchanged, mitigated but did not abolish the effect of the codon optimization on the CSC_g_ value of the fusion genes (Fig. S4B). If codon-optimization indeed specifically increased the mRNA half-life, the steady-state mRNA numbers should increase. Indeed, we found that the mean mRNA numbers increased for codon-optimized *mad2* and *mad3* compared to the wild-type gene (from 3.9 to 5.0 for *mad2*, and from 3.7 to 5.3 for *mad3*; Fig. 3D, S5C).

In *S. cerevisiae*, the RNA helicase Dhh1 (*S. pombe* Ste13) is involved in specifically lowering the mRNA half-life of genes with a high fraction of non-optimal codons (Cheng *et al*, 2017; Radhakrishnan *et al*, 2016). We therefore asked whether deletion of *ste13^+^* affects mRNA numbers or half-life of *mad2* and *mad3*. Indeed, deletion of *ste13^+^* increased the mRNA number but did not increase it further for codon-optimized *mad2* or *mad3* (Fig. 3D, S5C,D). Moreover, direct measurement of mRNA half-life showed that it was indeed prolonged for *mad2* and *mad3* mRNA in *ste13Δ* cells (from about 6 min to 10 min for *mad2*, and 4 min to 8 min for *mad3*; Fig. 3E, S5E). These results indicate that non-optimal codons in *mad2^+^* and *mad3^+^* contribute to the short mRNA half-life of these genes.

### Codon-optimization, but not *ste13^+^* deletion, increases the protein concentration of Mad2 and Mad3

To ask whether the consequences of codon-optimization propagate to the protein level, we quantified Mad2- and Mad3-GFP protein expressed from the wild-type or codon-optimized genes. Both immunoblotting and fluorescence microscopy showed an increase in protein concentration after codon-optimization (Fig. 4), presumably resulting from the increase in mRNA (Fig. 3), possibly enhanced by an increased translation efficiency. In contrast, the Mad2 and Mad3 protein concentrations in *ste13Δ* cells remained stable or even decreased when analyzed by immunoblotting (Fig. 4B,C). A plausible explanation for this result is that many mRNAs increase in concentration in *ste13Δ* cells and compete for translation resources such as ribosomes. Moreover, since the coding sequences of *mad2^+^* and *mad3^+^* remain unchanged, no specific enhancement in translation efficiency is expected. In summary, these data support that codon usage bias towards non-optimal codons in *mad2^+^* and *mad3^+^* supports their short mRNA half-life and thereby supports a gene expression pattern that lowers cell-to-cell variability.

**Figure 4.**
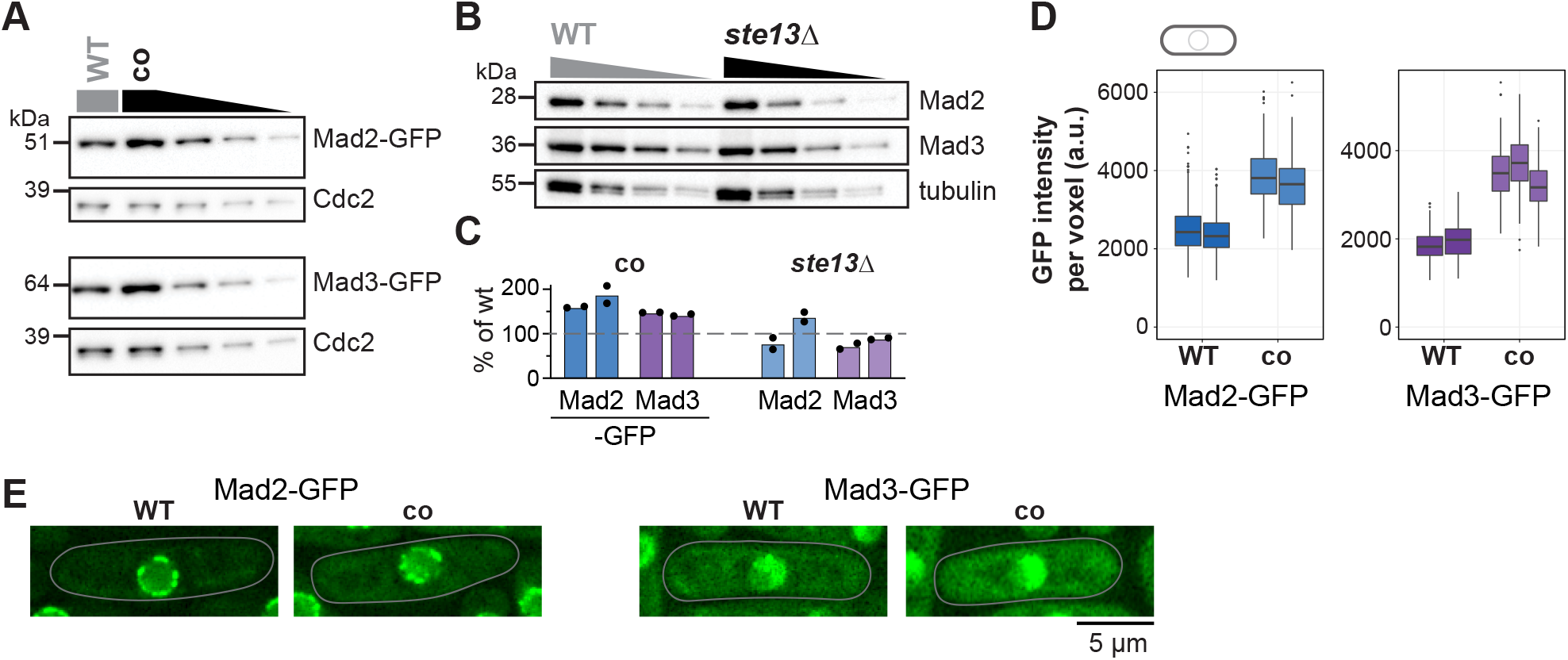
Codon optimization increases the protein concentrations of Mad2 and Mad3. **(A)** Immunoblot of protein extracts from cells expressing wild-type (WT) or codon-optimized (co) Mad2-GFP or Mad3-GFP probed with antibodies against GFP and Cdc2 (loading control). Lanes 3-5 are a 1:1 dilution series of the extract from cells expressing the codon-optimized version. **(B)** Immunoblot of protein extracts from wild-type (WT) or ste13Δ strains probed with antibodies against Mad2, Mad3 and tubulin (loading control). A 1:1 dilution series was loaded for quantification. **(C)** Estimates of the protein concentration relative to wild-type conditions from experiments such as in (A) and (B). Bars are biological replicates, dots are technical replicates. **(D)** Whole-cell GFP concentration from individual live cell fluorescence microscopy experiments (a.u. = arbitrary units). Boxplots show median and interquartile range (IQR); lines extend to values no further than 1.5 times the IQR from the first and third quartile, respectively. Mad2-GFP: n = 488 and 413; Mad2-co-GFP: n = 206 and 366; Mad3-GFP: n = 224 and 127; Mad3-co-GFP: n = 160, 450 and 212 cells. **(E)** Representative images from one of the experiments in (D). A single Z-slice is shown; cells are outlined in grey.

### *Mad1^+^* expression regulation differs from that of *mad2^+^* and *mad3^+^*

The *mad1^+^* gene shares a short mRNA half-life with *mad2^+^* and *mad3^+^* (Fig. 2D). Different from *mad2+* and *mad3+*, though, *mad1+* has a higher fraction of optimal codons and a CSC_g_ value above the median of all protein-coding *S. pombe* genes (Fig. 3A,B; S4A,B). This was surprising because we expected similar features within the SAC network. Consistent with its higher CSC_g_ value, the *mad1* mRNA numbers hardly changed after either codon-optimization or *ste13* deletion (Fig. 5A,B; S6A). After codon-optimization, the mRNA concentration (mRNA number per cell area) even decreased slightly (Fig. S6C,D). A second codon-optimized *mad1* sequence that was considerably different from the first (77% nucleotide identity in the coding sequence; Fig. S4F) showed the same effect (Fig. S6B,D). We still found that *mad1^+^* mRNA half-life was prolonged in *ste13Δ* cells (from 4 min to 7 min; Fig. 5C, S6E,F), but less so than for *mad2^+^* and *mad3^+^*. Thus, the short *mad1^+^* mRNA half-life is less dependent on codon usage bias. Notably, *mad2^+^* and *mad3^+^* mRNA, in addition to their codon non-optimality, contain motifs within their 3’ UTRs that have been associated with short mRNA half-life, whereas *mad1^+^* does not (Eser *et al.*, 2016). Hence, different modes of regulation bring about the short mRNA half-life of these SAC genes. The gene elements that contribute to the short mRNA half-life of *mad1^+^* remain to be identified.

**Figure 5.**
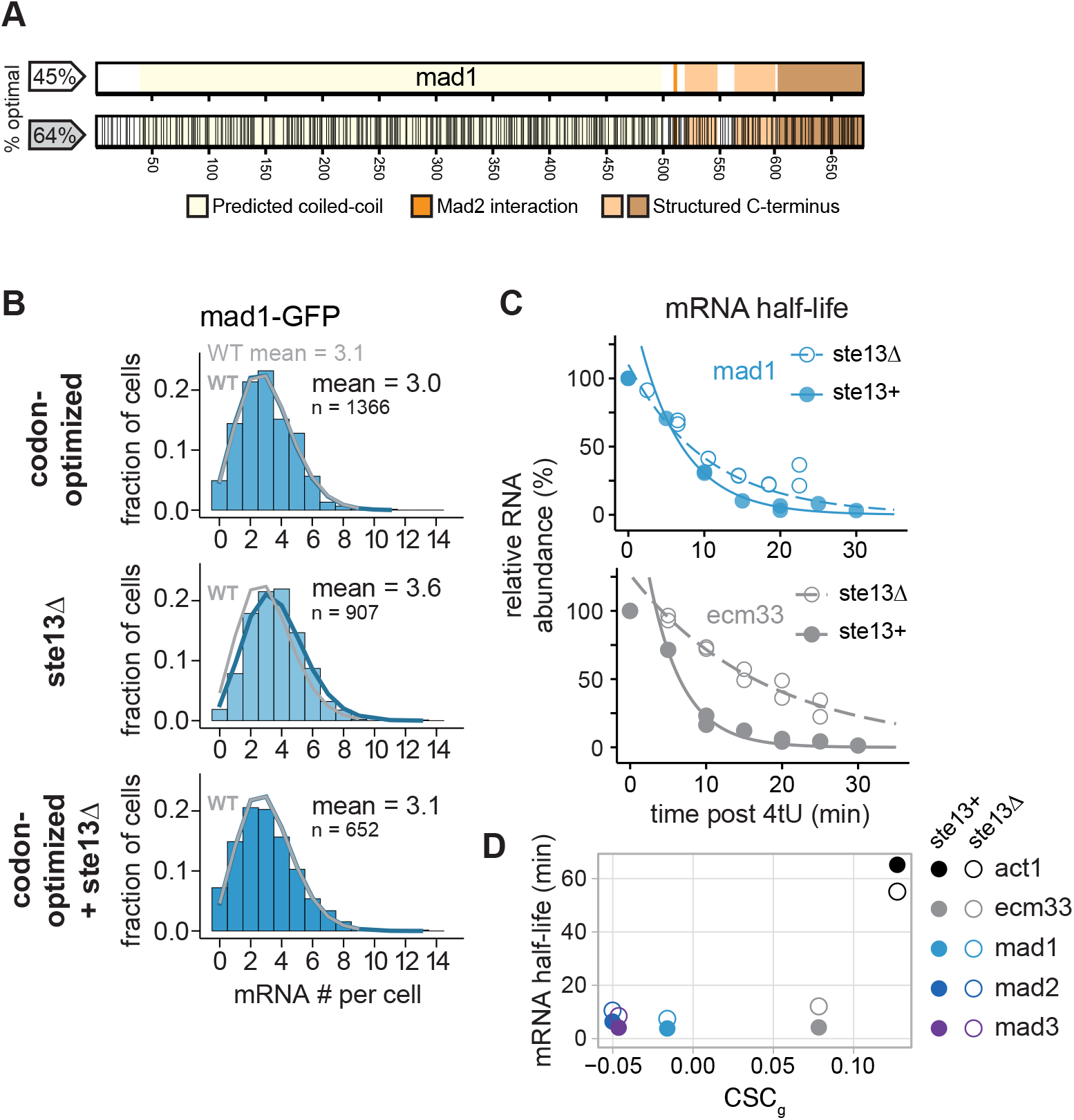
Codon optimization and *ste13* deletion do not greatly increase the steady-state mRNA number of *mad1*. **(A)** Schematic of the *mad1* gene. Regions coding for important structural features are highlighted. Black lines in the bottom graph indicate synonymous codon changes in the codon-optimized version. **(B)** Frequency distribution of mRNA numbers per cell determined by mRNA FISH. Probes were against the GFP portion of the *mad1* fusion gene. Curves show fit to a Poisson distribution. Grey curves show the fit from wild-type *mad1-GFP* cells (WT, Figure 1). Two replicate experiments were combined for codon-optimized genes and ste13 deletion (shown individually in Figure S4). **(C)** Time course of mRNA abundances by qPCR following metabolic labeling and removal of the labeled pool (two independent experiments). Lines indicate fit to one-phase exponential decay, excluding the measurements at t = 0 in order to accommodate for non-instantaneous labeling by 4tU. The *ste13^+^* data are the same as in Figure 2. **(D)** Comparison between mean CSC values for selected genes (CSC_g_) and mRNA half-life measured with or without deletion of *ste13*.

### Codon-optimization of *mad1^+^* decreases its protein concentration

Unlike Mad2- and Mad3-GFP, whose protein concentration increased after codon optimization, that of Mad1-GFP decreased, both by immunoblotting and fluorescence microscopy (Fig. 6A,C-E; S7A). Deletion of *ste13^+^* influenced Mad1-GFP protein concentration less (Fig. 6B,C). The lower protein concentration of Mad1 produced from codon-optimized mRNA may partly result from the slightly lower mRNA concentration (Fig. S6C,D). However, because Mad1 is a highly structured protein, it is also possible that a particular pattern of codon usage serves co-translational protein folding or co-translational complex formation, and this may have been disrupted by synonymous codon replacement (Liu *et al*, 2021). Problems in protein folding or complex formation may lead to co-translational degradation, or to production of a protein that is less stable (Walsh *et al*, 2020). Both could explain a lower protein concentration.

**Figure 6.**
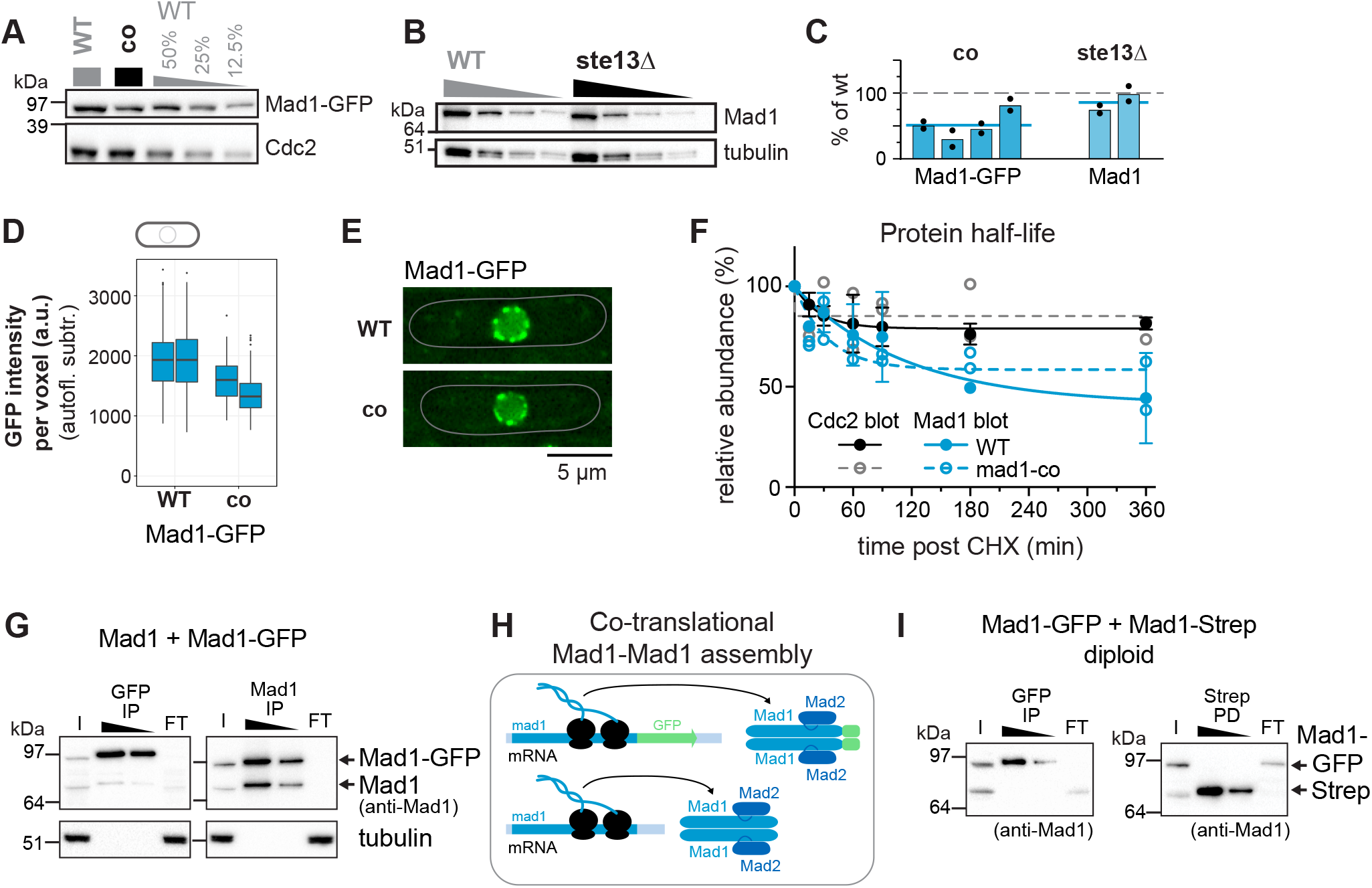
Codon identity in *mad1* is important for proper protein concentration. **(A)** Immunoblot of protein extracts from cells expressing wild-type (WT) or codon-optimized (co) Mad1-GFP probed with antibodies against GFP and Cdc2 (loading control). Lanes 3-5 are a dilution series of the extract from wild-type cells. **(B)** Immunoblot of protein extracts from wild-type (WT) or *ste13Δ* strains probed with antibodies against Mad1 and tubulin (loading control). A 1:1 dilution series was loaded for quantification. Tubulin blot is the same as in Figure 4. **(C)** Estimates of the protein concentration relative to wild-type conditions from experiments such as in (A) and (B). Bars are biological replicates, dots are technical replicates. Blue lines indicate the mean of all experiments. **(D)** Whole-cell GFP concentration from individual live cell fluorescence microscopy experiments (a.u. = arbitrary units). Boxplots show median and interquartile range (IQR); lines extend to values no further than 1.5 times the IQR from the first and third quartile, respectively. Mad1-GFP: n = 246 and 245; Mad1-co-GFP: n = 151 and 406 cells. **(E)** Representative images from one of the experiments in (D). An average projection of three Z-slices is shown; cells are outlined in grey. **(F)** Protein abundances after translation shut-off with cycloheximide (n = 3 experiments for wild-type (WT) cells, error bars = std; n = 2 experiments for mad1-co expressing cells). Lines indicate fit to a one-phase exponential decay. Extracts were probed for Mad1 and Cdc2. A representative immunoblot is shown in Figure S7A. **(G)** Immunoprecipitation (IP) with anti-GFP or anti-Mad1 from extracts of haploid cells expressing both untagged and GFP-tagged Mad1, probed with antibodies against Mad1 and tubulin. I, input; FT, flow-through. **(H)** Schematic illustrating that Mad1-Mad1 complex assembly likely takes place co-translationally, i.e. only proteins synthesized from the same mRNA are combined. **(I)** Anti-GFP immunoprecipitation (IP) and Strep pull-down (PD) from extracts of diploid cells expressing Mad1-GFP and Mad1-Strep from the two endogenous loci. The membrane was probed with antibodies against Mad1. I, input; FT, flow-through.

To assess protein stability, we shut off translation by cycloheximide. Protein levels dropped slightly faster for Mad1-GFP produced from codon-optimized mRNA compared to that produced from wild-type mRNA, but a large pool of Mad1-GFP persisted over several hours in both strains (Fig. 6F; S7B,C). This result is consistent with slight defects in folding caused by codon-optimization that led to some fraction of the protein becoming less stable.

Mad1 forms a homodimer through a long N-terminal coiled-coil, but whether this homodimer forms co-translationally or post-translationally was unknown. Co-translational formation may need a specific codon usage pattern. If homodimer formation was post-translational, it should be possible to observe interactions between tagged and untagged Mad1 that are expressed from two different loci in the same cell. However, in haploid strains expressing a C-terminally GFP-tagged and an untagged *mad1^+^* gene, a GFP immunoprecipitation almost exclusively precipitated Mad1-GFP, but not untagged Mad1 (Fig. 6G). In contrast, a Mad1 immunoprecipitation precipitated both Mad1-GFP and Mad1 equally. These experiments used a monomeric version of GFP. Thus, it is unlikely that this pattern is driven by dimerization of GFP and we instead suggest that Mad1 homodimers form co-translationally from the nascent chains of two ribosomes translating *mad1^+^* from the same mRNA molecule (Fig. 6H). We corroborated this finding by using diploid strains expressing Mad1-GFP and Mad1-Strep from the two endogenous loci. A GFP-immunoprecipitation isolated Mad1-GFP but not Mad1-Strep, whereas a Strep pull-down isolated Mad1-Strep but not Mad1-GFP (Fig. 6I). This preferential formation of homodimers was also seen for codon-optimized Mad1 (Fig. S7E). The fact that Mad1 homodimers form co-translationally is consistent with the idea that synonymous codon changes may subtly impair complex formation. Overall, these results suggest that codon usage bias within *mad1^+^* does not contribute to mRNA half-life shortening, but that codon usage within *mad1* may be optimized for production of correctly folded and dimerized protein.

### The coding sequence of *mad1^+^* influences mRNA expression

The slightly lower mRNA concentration after *mad1* codon-optimization suggested that the concentration of *mad1^+^* mRNA is not purely determined by regulatory sequences upstream and downstream of the coding sequence. This is supported by our observation that merely fusing GFP to *mad1^+^*, without altering surrounding sequences, increases its mRNA number (Fig. 1D).

Further supporting this notion, we found that replacing the *mad1^+^* coding sequence with GFP produced neither significant amounts of mRNA nor protein (Fig. 7). Hence, the sequences surrounding the *mad1^+^* coding sequence are insufficient to establish *mad1^+^*-like expression. Preserving the first 66 or 108 base pairs of *mad1^+^* partly rescued both mRNA and protein levels but not completely. While this suggests that the 5’ region of the *mad1^+^* coding sequence carries signals that are important for mRNA synthesis or stabilization, some other genes apparently contain sequences that can compensate. Introducing an *nmt1^+^-GFP* fusion gene (Fig. 7) or fusions between *S. cerevisiae* GCN4 and N-terminally truncated versions of *S. pombe mad1^+^* allowed for expression from the *mad1^+^* locus (Heinrich *et al*, 2014). What these genes share, that GFP does not, is unclear at present.

**Figure 7.**
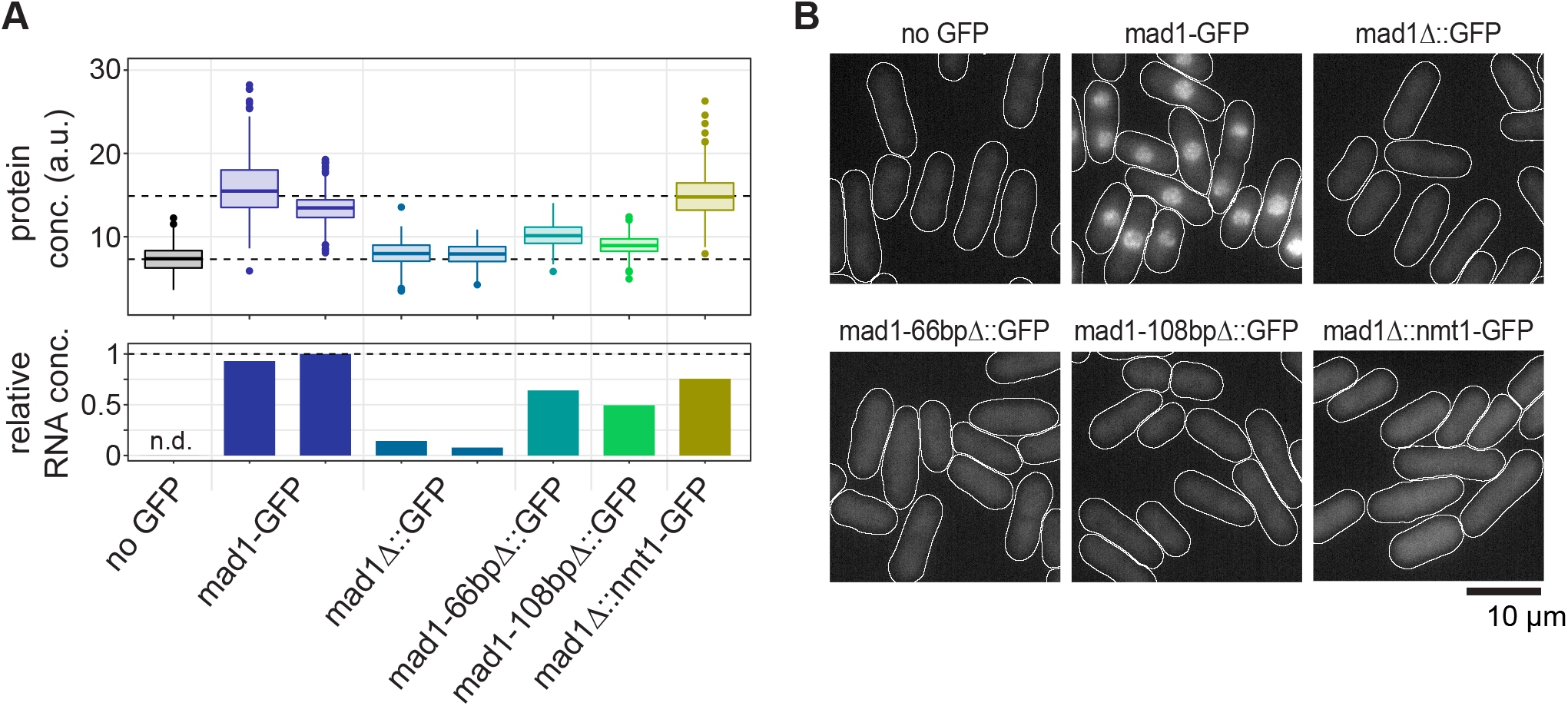
Sequences surrounding the *mad1* coding region are insufficient for proper expression. **(A)** The indicated strains were tested for GFP expression by qPCR (bottom panel) and by live-cell microscopy (top panel; a.u. = arbitrary units). *Mad1* was either fused to *GFP (mad1-GFP)*, or the coding sequence of *mad1* was replaced with *GFP (mad1Δ∷GFP*), with *nmt1-GFP (mad1Δ∷nmt1-GFP*) or with the 5’ region of *mad1* (66 bp or 108 bp) followed by *GFP*. A strain not expressing any GFP was used as reference. Boxplots show median and interquartile range (IQR); lines extend to values no further than 1.5 times the IQR from the first and third quartile, respectively. For microscopy: no GFP: n = 634; *mad1-GFP*: n = 525 and 391; *mad1Δ∷GFP*: n = 497 and 360; *mad1-66bpΔ∷GFP*: n = 293; *mad1-108bpΔ∷GFP*: n = 173; *mad1Δ∷nmt1-GFP*: n = 419 cells. **(B)** Representative live-cell microscopy images from the experiment in (A). Cell shapes are outlined in white.

Altogether, these results indicate that *mad1^+^* expression has some unique aspects: *mad1^+^* uses a different mode for reducing mRNA half-life than *mad2^+^* or *mad3^+^*, and its coding sequence carries elements that help transcribe or stabilize RNA.

## Discussion

Proteins are the workhorses of cells. The deployment of this workhorse army is controlled by regulatory elements encoded on DNA that are still incompletely understood. The spindle assembly checkpoint is sensitive to expression changes, and we therefore asked which features of gene expression may be important for its proper function. Our results suggest that a combination of short mRNA half-lives and long protein half-lives is important to keep protein variability low. We also find that—despite their closely shared function—*mad1^+^* differs in its expression features from *mad2^+^* and *mad3^+^*. The coding sequences of *mad2^+^* and *mad3^+^* contribute to the short mRNA half-life of these genes, whereas that of *mad1^+^* contributes to maintaining mRNA (Fig. 7) and protein levels (Fig. 6) in ways that we have so far been unable to resolve. We propose that the choice of synonymous codons in *mad1^+^* is optimized for the formation of the Mad1 homodimer and, ultimately, the Mad1/Mad2 complex.

### Short mRNA half-life of constitutively expressed SAC genes favours low noise

The short mRNA half-lives of *mad1^+^*, *mad2^+^* and *mad3^+^*, along with their long protein half-lives, can explain the low protein noise of SAC genes despite low and highly variable mRNA numbers (Fig. 1,2) (Thattai & van Oudenaarden, 2001). In human cells, a long protein half-life has also been shown to buffer the effects of variable mRNA numbers (Raj *et al*, 2006). Indeed, human Mad1, Mad2 and BubR1 (Mad3 ortholog) are also highly stable proteins (Rodriguez-Bravo *et al*, 2014; Schweizer *et al*, 2013; Suijkerbuijk *et al*, 2010; Varetti *et al*, 2011), which will support stable protein concentrations over time and between cells. SAC genes are probably not unique in combining a short mRNA and long protein half-life to achieve low noise. Other constitutively expressed genes that produce low or modest amounts of protein will likely show a similar behavior. Keeping noise low in this manner requires a high turn-over of mRNA that confers some energy cost. An alternative way to keep protein noise low would be to produce the same amount of protein from a larger number of more stable mRNA molecules (Fig. S3C). Several side-effects likely prohibit this solution as a general strategy. For example, the cytoplasm would be much more crowded with mRNAs, and stable mRNAs may accumulate chemical damage. Indeed, genes using an expression strategy of high transcription and low translation rates are exceedingly rare among different eukaryotes (Hausser *et al*, 2019).

### Different SAC genes employ different strategies for achieving short mRNA half-life

The half-life of an mRNA is influenced by sequence motifs, codon usage and other factors that influence translation. Currently, known factors predict around 50–60% of mRNA half-life in budding yeast (Cheng *et al.*, 2017; Neymotin *et al*, 2016). At least two elements seem to play a role for *mad2^+^* and *mad3^+^*: their mRNA half-lives are shortened by a high fraction of non-optimal codons (Fig. 3), and their 3’ UTRs contain sequence motifs that are associated with short mRNA half-life (Eser *et al.*, 2016). The predicted motifs in the 3’ UTR are likely functional because we found higher mRNA numbers after traditional tagging, which changed the 3’ UTR to that of a highly expressed gene (Heinrich *et al.*, 2013).

For *mad1^+^*, in contrast, the elements shortening its mRNA half-life remain unknown. Codon usage has no obvious influence (Fig. 5, S6), and the *mad1^+^* 3’ UTR does not seem to contain motifs implicated in half-life shortening (Eser *et al.*, 2016). We suspect that other elements that influence translation efficiency may play a role. Generally, less efficiently translated mRNAs are less stable (Hanson & Coller, 2018), and *mad1^+^* seems to be translated less efficiently than *mad2^+^* or *mad3^+^* (Rubio *et al*, 2020).

While our results show that *S. pombe* Ste13—similar to *S.c.* Dhh1—can indeed shorten mRNA half-life (Fig. 3, 5), it remains unclear what makes an mRNA a strong target of Ste13. We were surprised to find that our control gene *ecm33+* is an excellent target despite its high fraction of optimal codons (Fig. 5C,D; S4A, S6G) and high translation efficiency (Rubio *et al.*, 2020). Overall, our results support known elements of regulation but also highlight current gaps in understanding.

### Formation of the Mad1/Mad2 complex involves co-translation assembly of the Mad1 dimer but not synchronous assembly of the tetramer

Mad1 and Mad2 form a tight tetrameric complex (Kim *et al.*, 2012; Sironi *et al.*, 2002), but how this complex assembles is unknown. Our experiments suggest that the Mad1 homodimer forms between two polypeptides translated from the same mRNA, and that Mad1 molecules translated from different mRNA molecules associate very inefficiently with each other, if at all (Fig. 6). This assembly mode is further supported by a recent proteome-wide analysis in human cells that came to the same conclusion for Mad1 and other proteins with N-terminal dimerization through a coiled-coil (Bertolini *et al*, 2021). At least two studies have expressed Mad1 N-terminal fragments and full-length Mad1 from two different loci and have interpreted the failure to see association between those two as an inability of the N-terminal fragment to dimerize (Ji *et al*, 2018; Jin *et al*, 1998). However, we suggest that the capacity of an N-terminal Mad1 fragment to dimerize would need to be based on assessing self-association rather than assessing association with Mad1 expressed from a different locus. Of note, C-terminal Mad1 fragments also dimerize, but this dimerization may be able to occur post-translationally (Kim *et al.*, 2012).

While we suggest that assembly of the Mad1 homodimer occurs co-translationally, the assembly of the Mad1/Mad2 tetramer does not occur in synchronous co-translational fashion, since the mRNAs for *mad1^+^* and *mad2^+^* do not co-localize in the cytoplasm (Fig. 1). This leaves open the possibility of post-translational assembly of the tetramer or of asynchronous co-translational assembly, where one protein is already fully formed and binds the other that is being translated (Duncan & Mata, 2011; Shiber *et al*, 2018). Formation of the C-Mad2/Cdc20 complex necessitates catalysis (Faesen *et al*, 2017; Kulukian *et al*, 2009; Lad *et al*, 2009; Piano *et al.*, 2021; Simonetta *et al*, 2009), making it likely that C-Mad2/Mad1 formation also needs to be facilitated. We favour the idea that the tetramer assembles while one of the proteins is being translated, and it will be interesting to test whether the *mad1^+^* mRNA binds Mad2 protein or vice versa to facilitate such an assembly.

### Potential SAC malfunction from synonymous mutations

Overall, our data suggest that the coding sequences of *mad1^+^*, *mad2^+^* and *mad3^+^* modulate gene expression. Hence, even synonymous mutations carry some risk of impairing the SAC. We suspect that *mad1* is most susceptible to single synonymous substitutions, given the strong influence of the 5’ end of the coding sequence on transcription (Fig. 7) and the need for co-translational homodimer assembly (Fig. 6), which may be facilitated by controlling the speed of ribosome movement. For example, a cluster of non-optimal codons follows the coiled-coil region in *S. pombe mad1^+^* (Fig. S4E, S8), which may ensure that the N-terminal coiled-coil is fully formed before the remainder of Mad1 is translated.

It will be interesting to test whether synonymous mutations found in cancer samples can modulate SAC gene expression or function. Within MAD2L1 (*H.s. mad2*), synonymous mutations detected in cancer samples seem to cluster in a conserved region with high CSC values preceding the ‘seat belt’ (Fig. S8), suggesting that codon usage bias in this region may be functionally important. Although most synonymous mutations will only have small effects, they may fuel carcinogenesis. This is particularly true in the context of the SAC, because drastic impairment is more likely to be detrimental for cancer cells whereas subtle impairment may promote carcinogenesis (Cohen-Sharir *et al.*, 2021; Funk *et al.*, 2016; Kops *et al.*, 2004; Quinton *et al.*, 2021). The role of synonymous mutations and changes in tRNA expression in cancer is more and more recognized (Sauna & Kimchi-Sarfaty, 2011; Supek *et al*, 2014), and our data suggest that the SAC may be no exception.

## Materials and Methods

### Yeast strains

Yeast strains are listed in Table S1. Tagging of *nmt1^+^* and deletion of *ste13^+^* was done by conventional PCR-based gene targeting (Bähler *et al*, 1998). Marker-less insertion at the endogenous locus was performed using CRISPR/Cas9 (Jacobs *et al.*, 2014). Sequences used for targeting Cas9 to different loci are listed in Table S2. The *mad2^+^-ymEGFP* strain contains a single, silent (AGG to AGA) PAM site mutation at amino acid position 173 of Mad2. The *mad3^+^-ymEGFP* strain contains a single, silent (TTG to TTA) PAM site mutation at amino acid position 199 of Mad3. Yeast, monomeric enhanced GFP (ymEGFP) was derived from yEGFP (yeast codon optimized green fluorescent protein (Watson *et al*, 2008)) by mutation of Alanine 206 to Arginine (A206R), which is expected to reduce dimerization (Zacharias *et al*, 2002). Codon-optimization used proprietary algorithms by two different companies, and sequences are listed in Table S3. The strain with two differently tagged versions of *mad1^+^* has *mad1^+^*-*ymEGFP* along with 110 bp upstream and 164 bp downstream of the coding sequence integrated between the *leu1^+^* and *apc10^+^* gene.

### Yeast cultures

*S. pombe* cultures were grown at 30 °C either in rich medium (yeast extract supplemented with 0.15 g/L adenine; YEA) or in Edinburgh minimal medium (EMM, MP Biomedicals, 4110012) supplemented with 0.2 g/L leucine, 0.15 g/L adenine or 0.05 g/L uracil if required (Petersen & Russell, 2016). When cultures in minimal medium were started at low concentration, ‘pre-conditioned medium’ was added to a maximum of 50%. Pre-conditioned medium was obtained by growing cells in EMM and then removing the cells by centrifugation and filtration. For growth assays, cells were grown in YEA to a concentration of around 1 × 10^7^ cells/mL, diluted to 4 × 10^5^ cells/mL in YEA and further diluted in a 1:5 series of dilutions. 10 *μ*L were spotted on indicated plates. *S. cerevisiae* cultures were grown at 30 °C in bacto yeast extract supplemented with 20 mg/mL each of bacto peptone and dextrose (YPD).

### Cycloheximide treatment for determination of protein half-lives

Cells were grown in EMM (plus supplements required for auxotrophic mutations) to a final concentration of around 1 × 10^7^ cells/mL. Cultures were diluted to 8 × 10^6^ cells/mL, transferred to a 30 °C water bath for 30 minutes and a sample was taken prior to addition of cycloheximide (CHX) to a final concentration of 1 mg/mL. Cells were collected at specified timepoints, spun down at 980 rcf and frozen in liquid nitrogen before processing.

### TCA whole cell extracts

Cells were grown to a final concentration of around 1 × 10^7^ cells/mL and collected by centrifugation. Supernatant was removed and cells were washed with 1 mL of 20% trichloroacetic acid (TCA). Supernatant was removed and cells were resuspended in 500 *μ*L of water. 75 *μ*L of NaOH/beta-mercaptoethanol (final conc. = 0.22 M NaOH, 0.12 M b-ME) was added and samples incubated on ice for 15 minutes. 75 *μ*L of 55% TCA was added and samples incubated on ice for another 10 minutes. Samples were spun at 16,900 rcf for 10 minutes at 4 °C, and supernatant was removed. Pellets were resuspended in 100 *μ*L sample buffer (50 *μ*L of 2x HU buffer [8 M urea, 5% SDS (w/v), 200 mM Tris-HCl pH 6.8 (v/v), 20% glycerol (v/v), 1 mM EDTA (v/v), 0.1% (w/v) bromophenol blue], 40 *μ*L water and 10 *μ*L of 1 M DTT) to a final concentration corresponding to 1 × 10^9^ cells/mL. Approximately 150 *μ*L of acid washed beads (Sigma) were added before agitation in a ball mill (Mixer Mill 400; Retsch) for 2 minutes at 30 Hz. Tubes were pierced at the bottom, cell extract was collected from the beads by centrifugation and heated at 75 °C for 5 minutes.

### Immunoblotting

Proteins were separated by SDS-PAGE (NuPAGE, Bis-Tris, Thermo Fisher) and transferred onto PVDF membranes (Immobilon-P, Millipore) using a semi-dry blotting assembly (Amersham Biosciences TE-70 ECL semi-dry unit) and transfer buffer with 10% methanol. Membranes were probed with mouse anti-GFP (Roche, 11814460001), rabbit anti-Cdc2 (CDK1, Santa Cruz, SC-53), mouse anti-Cdc13 (cyclin B, Novus, NB200-576), rabbit anti-Mad1 (Heinrich *et al.*, 2013), rabbit anti-Mad2 (Heinrich *et al.*, 2013), rabbit anti-Mad3 (Heinrich *et al.*, 2013) or mouse anti-tubulin (Sigma, T5168). Secondary antibodies were either anti-mouse or anti-rabbit conjugated to HRP (Dianova) and quantified by chemiluminescence using SuperSignal West Dura ECL (ThermoFisher) and imaged on a Bio-Rad Gel Doc system. Chemiluminescence signals were quantified on non-saturated images using Image Lab software (Bio-Rad). Measurements from a reference dilution series were used to create a standard curve, which was used to determine the concentration of sample relative to the reference.

### Immunoprecipitation

Asynchronously growing cultures were harvested, washed with deionized water and frozen as droplets in liquid nitrogen. Cell powder was prepared from these droplets using a ball mill (Mixer Mill 400; Retsch) for 30 seconds at 30 Hz under cryogenic conditions. Cell powder was resuspended in lysis buffer (20 mM Tris pH 7.5, 150 mM NaCl, 5% glycerol, 0.1% NP-40, 5 mM MgCl2) and protein concentration was determined by BCA assay (ThermoFisher). For immunoprecipitation, powder was resuspended to a final concentration of 20 mg/mL in lysis buffer supplemented with a 10x final concentration of protease (ThermoFisher, 87785) and phosphatase inhibitor cocktails (ThermoFisher, 78420). Extracts were spun down for 10 minutes at 4 °C and 16,900 rcf. As input sample, supernatant was mixed with an equal volume of sample buffer (2x HU buffer with 200 mM DTT) and heated for 3 minutes at 75 °C. For immunoprecipitations, the remaining extract was mixed with 50 *μ*L of Protein G Dynabeads (ThermoFisher, 10004D) coupled with anti-GFP antibodies (Roche, 11814460001) or anti-Mad1 antibodies (Heinrich *et al.*, 2013) and incubated with gentle agitation at 4 °C for 10 minutes. Supernatant was retained (flow through), and beads were washed three times with lysis buffer, gently agitated for 2 minutes at 4 °C between washing steps. Flow through was mixed with an equal volume of sample buffer and heated for 3 minutes at 75 °C. Elution from beads was performed by addition of 7 *μ*L 100 mM citric acid and gentle agitation for 5 minutes at 4 °C. Samples were neutralized by addition of 1.4 *μ*L of 1.5 M Tris pH 9.2, mixed with an equal volume of sample buffer and heated at 75 °C for 3 minutes. The Streptag pulldown was completed in the same manner using 100 *μ*L of MagStrep "type3" XT bead suspension (IBA Lifesciences, 2-4090-002), except elution was achieved by boiling with sample buffer at 95 °C for 2 minutes.

### Quantification of GFP fusion proteins in single cells (3D segmentation)

To quantify GFP fusion proteins in single cells, cells were grown in EMM (plus supplements that were required for auxotrophic mutations) at 30 °C to a final concentration of 6–9 × 10^6^ cells/mL. Cultures of GFP-positive and GFP-negative cells were mixed at a 1:1 ratio to a final concentration of 2.5–6.0 × 10^6^ cells/mL and incubated for 30 minutes at 30 °C. To ensure a uniform and flat imaging plane, cells were loaded into a Y04C microfluidics trapping plate (Millipore Sigma) and incubated inside a climate-controlled microscope chamber for 2 hours at 30 °C with constant flow of fresh media. Imaging was performed on a DeltaVision Elite system equipped with a PCO edge sCMOS camera and an Olympus 60x/1.42 Plan APO oil objective. Images were acquired for ymEGFP, tdTomato and brightfield as 7.2 *μ*m or 10 *μ*m stacks with images separated by 0.1 *μ*m. The acquired image area was 1046 × 1046 pixels with 1 × 1 binning. All images were deconvolved using SoftWoRx software. To correct for uneven illumination, deconvolved fluorescence images were flatfielded individually for each channel using a custom FIJI script (Baybay *et al.*, 2020).

The Pomegranate image analysis pipeline (Baybay *et al.*, 2020) was used to segment nuclei (using TetR-tdTomato-NLS) and whole cells (using brightfield signal and spherical extrusion of the midplane segmentation) (Fig. S2A). We corrected for chromatic aberration and for stretching of distances in the Z direction (Baybay *et al.*, 2020). Further analysis was conducted in R (R-Core-Team, 2020) and figures were produced using the package ggplot2 (Wickham, 2016).

Only information from mono-nucleated cells for which both the whole cell and the nucleus had been segmented was retained. Cells were excluded if one or more of the following conditions were met: the nuclear segmentation protruded beyond the three-dimensional bounds of the cell; whole-cell segmentation was cut-off by more than two slices because insufficient slices in Z had been recorded; cell was at the image edge and incompletely recorded; the nucleus had an aspect ratio (diameter in Z to diameter in XY) of less than 0.8 or more than 1.2; cell volume was in the 0.1^st^ or 99.9^th^ percentile. Cells with or without GFP signal were distinguished by k-means (k = 2) clustering (Fig. S2D-F), except for Nmt1-GFP, where the threshold for each image was set manually. One image, where the autofluorescence of GFP-negative cells deviated by more than three standard deviations from that of other images, was excluded. One additional image, where the cells had visibly moved during acquisition, was also excluded.

To subtract autofluorescence and other background, we averaged the fluorescence intensity per cell or nuclear volume for GFP-negative cells in an image and subtracted that value from the fluorescence intensity per cell or nuclear volume of each GFP-positive cell in the image. For a rough estimate of absolute concentration in nanomolar, we used our previous estimate of about 70 nM Mad3-GFP in the cell nucleus (Heinrich *et al.*, 2013) and normalized all background-subtracted data to this value.

Even after background subtraction, we observed some variation of mean intensities between single images (Fig. S2F) and we could not distinguish whether these differences were a consequence of sampling or came from conditions on the microscope stage while recording the image. We therefore opted to determine the coefficient of variation (CV = standard deviation / mean) for each protein not across all images, but instead for each image separately; Fig. 2A shows the variation across images.

### Quantification of GFP signals in single cells (2D segmentation and projection)

For experiments evaluating fluorescence signals after replacing the coding sequence of *mad1^+^* (Fig. 7), quantification was performed on projections, using 2D segmentation of cells. Cells were grown in minimal medium, collected by centrifugation from liquid cultures, mounted in medium on a slide, and brightfield and fluorescence images were collected immediately at room temperature. At least two slides were prepared and imaged for each strain. Images were recorded on a Zeiss AxioImager M1, using Xcite Fire LED illumination (Excelitas), a Zeiss Plan-Apochromat 63x/1.40 Oil DIC objective and an ORCA-Flash4.0LT sCMOS camera (Hamamatsu) with Z sections spaced by 0.2 *μ*m.

Cells were segmented based on an in-focus brightfield image using YeaZ (Dietler *et al*, 2020). Falsely segmented cells (e.g. background, or cells falsely combined into one) were manually excluded in Fiji. Only cells in the center of the image, where fluorescence illumination was homogeneous, were included. Flatfielding was not performed. The brightfield images were systematically shifted relative to the fluorescence images and we corrected for that error. Quantification of signals was performed on an average projection of the 23 most in-focus Z-slices (covering 4.6 *μ*m, which is slightly larger than the width of a typical *S. pombe* cell). For each image, the median extracellular background in the same central area of the image was subtracted.

### Single-molecule mRNA FISH

For quantification of mRNA by single-molecule fluorescent in-situ hybridization (smFISH), cultures of asynchronously dividing cells were grown to a concentration of about 1 × 10^7^ cells/mL in EMM. Typically, 2 × 10^8^ cells were fixed with 4% paraformaldehyde for 30 minutes before being washed three times with ice-cold Buffer B (1.2 M sorbitol, 100 mM potassium phosphate buffer pH 7.5) and stored at 4 °C before digestion of the cell wall. Cells were resuspended in spheroblast buffer (1.2 M sorbitol, 0.1 M potassium phosphate, 20 mM vanadyl ribonuclease complex [NEB S1402S], 20 *μ*M beta-mercaptoethanol) and digested with 0.002% 100T zymolyase (US Biological Z1005) for approximately 45–75 minutes. Zymolyase reaction was quenched when addition of water to the cells resulted in around 50% lysed cells. Reactions were quenched with 3 washes of Buffer B. Cell pellets were resuspended in 1 mL of 0.01% Triton X-100 in 1x PBS for 20 minutes and washed three times with Buffer B. For hybridization of probes, approximately 20–25 ng of CAL Fluor red 610 probes targeting ymEGFP or *mad2*, or Quasar 570 probes targeting *mad1* were mixed with 2 *μ*L each of yeast tRNA (Life Technologies) and Salmon sperm DNA (Life Technologies) per reaction. For two-color FISH experiments, 20–25 ng of each probe were combined, resulting in ~50 ng of total FISH probes per reaction. Sequences of probes are given in Table S4. Buffer F (20% formamide, 10 mM sodium phosphate buffer pH 7.2; 45 *μ*L per reaction) was mixed with the probe solution, heated at 95 °C for 3 minutes and allowed to cool to room temperature before mixing with Buffer H (4x saline-sodium citrate (SSC) buffer, 4 mg/mL acetylated BSA, 20 mM vanadyl ribonuclease complex; 50 *μ*L per reaction). Each sample of digested cells was divided into two reactions, each of which was resuspended in 100 *μ*L of this hybridization solution. Resuspended cells were incubated at 37 °C overnight. Cells were washed with 10% formamide/2x SSC followed by 0.1% Triton X-100/2x SSC). For DAPI staining, cells were incubated in 1x PBS with 1 *μ*g/mL DAPI for 10 minutes and washed once more with 1x PBS. Cell pellets were mixed with SlowFade Diamond Antifade Mountant (Thermo Scientific, S36972) and mounted on DEPC-cleaned slides using #1 1/2 glass coverslips. Imaging was performed on a Zeiss AxioImager M1 equipped with Xcite Fire LED illumination (Excelitas), a Zeiss *α* Plan FLUAR 100x/1.45 oil objective and an ORCA-Flash4.0LT sCMOS camera (Hamamatsu). Images were acquired for 6 *μ*m in Z separated by 0.2 *μ*m steps for each channel. Images of labelled RNA were captured with either an mCherry filter or a ‘gold FISH’ filter (Chroma, 49304). Additional data on the cell and nucleus were captured with GFP, DAPI and CFP. Images were background subtracted and flatfield corrected. A custom FIJI macro, using trainable WEKA segmentation (Arganda-Carreras *et al*, 2017), was used to create the outlines of cells by CFP autofluorescence and of corresponding nuclei by DAPI. RNA spot analysis was performed in FISHquant (Mueller *et al*, 2013). Spots were initially detected based on an automatic intensity threshold and filtered with an additional manual threshold following the suggestions of the FISHquant documentation. A subset of cells in each image was cross-checked manually for successful RNA spot detection. The frequency distribution of total and cytoplasmic mRNA per cell was approximately Poisson and fit with a Poisson distribution. FISHquant output was processed in R (R-Core-Team, 2020) using packages tidyverse (Wickham *et al*, 2019), broom, nabor, spatstat (Baddeley *et al*, 2016), mclust (Scrucca *et al*, 2016) and plyr (Wickham, 2011).

To measure co-localization of *mad1* and *mad2* mRNA, a two-color FISH experiment was performed targeting *mad1* with gene-specific probes and *mad2^+^-ymEGFP* with ymEGFP probes. The three-dimensional coordinates of each spot were recorded and corrected for relative chromatic aberration in Z. Distances were then calculated from each mRNA to its nearest neighbor of the other species within the same cell. In order to determine a distance cutoff to use for classifying RNA molecules as either co-localized or unpaired, the same two probe sets were used in another two-color FISH experiment in which both probes targeted *mad1^+^-ymEGFP*. Nearest-neighbor distances were calculated in the same way, and the distribution of these distances was used to determine the co-localization distance cutoff value. This cutoff was applied to the distances in the original experiment to classify each *mad1* or *mad2* mRNA molecule as co-localized or unpaired.

### Quantification of Mad1-GFP signals at kinetochores

Strains expressing the tubulin mutant *nda3-KM311* were grown in EMM (plus supplements required for auxotrophic mutations) at 30 °C to a concentration between 0.5–1.0 × 10^7^ cells/mL. Cells were diluted with EMM to a final concentration of 7.5 × 10^5^ or 1.5 × 10^6^ cells/mL. 300 *μ*L of each strain was loaded into a lectin-coated Ibidi *μ*-Slide glass-bottom chamber and incubated about one hour at 16 °C on the microscope stage prior to imaging. Cells were imaged at 16 °C on a DeltaVision Elite system with a PCO edge sCMOS camera (PCO) and an Olympus 60x/1.42 Plan APO oil objective and EMBL environmental chamber. Images were acquired every 5 minutes for GFP and mCherry over an 18-hour period using an ‘optical axis integration’ (sum projection) over a 3.2 *μ*m Z-distance.

Plo1-mCherry localizes to spindle pole bodies during mitosis and was used to identify cells entering mitosis. Kinetochores cluster with spindle pole bodies in *S. pombe* interphase (Funabiki *et al*, 1993) and dot-like GFP signals were therefore measured in the direct vicinity to Plo1-mCherry. An area of the same size for each cell was used to capture the kinetochore signal and was also used to measure the intensity in the nucleoplasm for background subtraction. GFP intensities from multiple cells were aligned to the time point of Plo1-mCherry appearance and averaged for each timepoint.

### RNA Preparation

Asynchronous *S. pombe* cultures were grown to a final concentration of approximately 0.7–1.5 × 10^7^ cells/mL at 30 °C in either EMM with 0.2 g/L leucine or YEA. 1 × 10^8^ cells were collected by centrifugation, washed once with deionized water, and immediately flash-frozen in liquid nitrogen and stored at −80 °C before processing. RNA was extracted by resuspending samples in 700 *μ*L of ice-cold TES buffer (10 mM Tris HCL pH 7.5, 10 mM EDTA, 0.5% SDS) and adding 700 *μ*L of acidic phenol chloroform (Fisher Scientific). Samples were immediately vortexed for 20 seconds and incubated for 1 hour at 65 °C. Following incubation, samples were cooled on ice for 1 min, vortexed for an additional 20 seconds and centrifuged for 15 minutes at 16,000 rcf at 4 °C. The RNA was further purified by twice mixing the aqueous supernatant with 700 *μ*L of acidic phenol chloroform and centrifuging the solution in a 5Prime Phase Lock Gel Heavy 2 ml tube (Andwin Scientific) at 16,000 rcf to separate the phases. Following overnight ethanol precipitation, samples were centrifuged at 16,000 rcf for 10 minutes at 4 °C and washed with one equivalent of 70% ethanol before additional centrifugation. Samples were left to air dry at room temperature and resuspended in nuclease free water before quantification. 50 *μ*g of total RNA was subjected to DNase treatment (Roche, 10 776 785 001) followed by ethanol precipitation.

### Quantitative PCR (qPCR)

For quantitative PCR (qPCR), 1 *μ*g of DNase-treated total RNA was subjected to Superscript IV cDNA synthesis using oligo d(T)_20_ primers. Transcript abundance was quantified on a QuantStudio 6 Real Time PCR system using SYBR® Green PCR Master Mix (ThermoFisher) and gene-specific primers (Supplementary Table S5). To estimate relative expression, raw Ct values (2–3 technical replicates per sample) were averaged and normalized according to the following formula (Hellemans *et al*, 2007; Pfaffl, 2001):

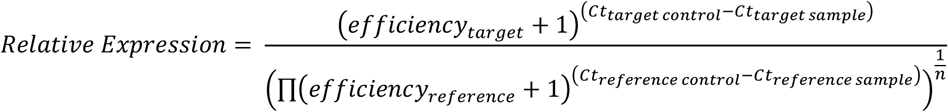

Where “target” is the mRNA of interest, “reference” is the reference gene, “sample” is the sample of interest and “control” is the control sample being normalized to. The denominator is the geometric mean of the reference genes (*act1* and *cdc2*), and efficiencies were estimated from the slopes of four-step, serial 1:5 dilution standard curves.

### Determination of mRNA half-life

The mRNA half-life measurement protocol was adapted from Duffy et al. 2015 (Duffy *et al*, 2015) and Chan et al. 2018 (Chan *et al*, 2018). Asynchronous *S. pombe* cultures were grown to a final density of approximately 0.7–0.9 × 10^7^ cells/mL at 30 °C in EMM with 0.45 mM uracil and 1.5 mM leucine before collection. 4TU in DMSO (Chem-Impex International) was added at 5 mM final concentration (Eser *et al.*, 2016). Cells (5 × 10^7^ cells per sample) were collected by centrifugation, and immediately flash-frozen in liquid nitrogen and stored at −80 °C before processing. Samples were collected from the culture before addition of 4TU (time = 0) and at a series of time points after. For use as a spike-in control, *S. cerevisiae* cultures were grown to a final density of 1.4–3.7 × 10^7^ cells/mL in YPD at 30 °C, and then flash-frozen and stored at −80 °C.

RNA extraction was performed as above except flash-frozen samples were initially resuspended in 600 *μ*L of ice-cold TES buffer and 100 *μ*L of resuspended *S. cerevisiae* cells (approximately 5 × 10^7^ cells) were added as a spike-in control. Care was taken to add the same amount of *S. cerevisiae* cells to all sample of a time series. After extraction, 200 *μ*g of total RNA was subjected to DNase treatment. Following DNase treatment, 70 *μ*g of RNA was biotinylated with MTSEA biotin-XX (Biotium) as previously described (Chan *et al.*, 2018; Duffy *et al.*, 2015). 50 *μ*g of biotinylated RNA was subjected to oligo d(T) selection using oligo d(T)_25_ magnetic beads (NEB S1419S), substituting SDS and NaCl for the recommended LiDS and LiCl. For streptavidin selection of biotinylated RNA, 500 ng of the oligo d(T) selected mRNA was used. 25 *μ*L of MyOne Streptavidin C1 Dynabeads (ThermoFisher 65001) were washed with 75 *μ*L of 0.1 M NaOH two times, followed by a single 0.1 M NaCl wash, and two additional washes with Buffer 3 (10 mM Tris HCl pH 7.4, 10 mM EDTA, 1 M NaCl). Streptavidin beads were blocked by resuspension in 50 *μ*L of Buffer 3 and 5 *μ*L of 50x Denhardt’s reagent. Beads were incubated for 20 minutes with gentle agitation. Following blocking, beads were washed with 75 *μ*L of Buffer 3 four times and resuspended in 75 *μ*L of Buffer 3 with 4 *μ*L of 5 M NaCl. 500 ng of mRNA was added to the beads and gently agitated for 15 minutes. Following incubation, beads were washed with 75 *μ*L of Buffer 3 prewarmed to 65 °C, once with Buffer 4 (10 mM Tris-HCl pH 7.4, 1 mM EDTA, 1% SDS) and twice with 10% Buffer 3. All flow through was pooled before addition of 20 *μ*g of linear acrylamide (ThermoFisher) followed by ethanol precipitation.

Expression relative to the time = 0 sample was quantified using qPCR as described above except 50 ng of recovered unlabeled mRNA was used in the SSIV cDNA synthesis reaction and a single reference gene (*S. cerevisiae ACT1* from the spiked in *S. cerevisiae* cells) was used to normalize expression.

To estimate mRNA half-lives (*HL*), several different exponential decay models (adapted from (Chan *et al.*, 2018)) were fit to each time series using non-linear least squares regression:

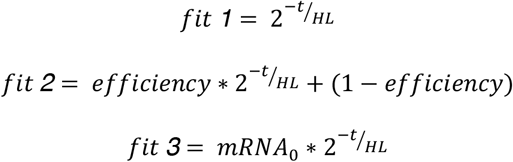

*Fit 1* is a simple one-phase exponential decay model. *Fit 2* incorporates an efficiency parameter to accommodate that mRNA levels may not decay to zero (Chan *et al.*, 2018). Finally, to accommodate the effects of non-instantaneous labelling by 4TU on the decay curve, the *fit 3* model was fit without the time = 0 measurements and instead allowed the mRNA level at time = 0 to be estimated as a separate free parameter (*mRNA_0_*). Model fitting was done in R using the function nls() from the package “stats” (R-Core-Team, 2020). Qualitatively, *fit 3* consistently fit the time series the best, and half-lives estimated with this model are presented in the text. Half-life estimates from all three models are presented in Figs. S5E and S6F.

### Codon usage bias calculations

The “Codon occurrence to mRNA Stability Correlation coefficient” (CSC) for each codon was determined as in Presnyak et al. (Presnyak *et al.*, 2015) by calculating a Pearson correlation coefficient between the frequency of occurrence of individual codons in mRNAs and the half-lives of these mRNAs. Coding sequences for *S. pombe* (protein-coding genes, excluding dubious and transposons) were downloaded from Pombase (ASM294v2.62, Release date 2017-01-30) (Lock *et al*, 2019). From this list we excluded “Genome location: mitochondrial”, “Genome location: mating_type_region” and “sequence error in genomic data” (PBO:0000129). Five genes lacking start or stop codons were additionally excluded, resulting in a final list of 5,016 genes. We used mRNA half-life data from either Hasan et al. (Hasan *et al.*, 2014) or Eser et al. (Eser *et al.*, 2016), which are the most recent and comprehensive datasets for *S. pombe*. Out of the 5,016 genes in our list, 4,615 were measured in at least one study and 3,900 in both. Both studies used metabolic labelling and the half-lives correlate with each other (Pearson correlation coefficient 0.50; Spearman’s rank correlation 0.81).

A previous study (Harigaya & Parker, 2016) had used the Spearman correlation coefficient to determine CSC values for *S. pombe*, because of outliers in the half-life data. We instead removed outliers from the half-life data, and then used the Pearson correlation coefficient. A comparison between the different strategies is shown in Fig. S4C,D. Our criteria for removing outliers were: (i) a value that was more than 10 interquartile ranges above the upper quartile (which removed three genes, based on their value in the Eser et al. data) and (ii) a deviation in rank position of > 2,500 between the two datasets (which removed 13 genes). After the removal of outliers, the Pearson correlation coefficient between the two mRNA half-life datasets was 0.80, the Spearman’s rank correlation 0.82. Using either the Hasan et al. or the Eser et al. half-life data for CSC calculation yielded highly similar results (Fig. S4C,D). When not otherwise indicated, CSC values obtained from Eser et al. (the more recent study) were used. The CSC_g_ value for each gene was determined as the arithmetic mean of all codons, excluding the stop codon.

The percentage of optimal codons based on the “classical translational efficiency” (cTE) used the optimality table for *S. pombe* from Pechmann and Frydman (Pechmann & Frydman, 2013). For tAI (tRNA adaptation index), we used the values determined by Tuller et al. (Tuller *et al*, 2010) and also reported in Pechmann and Frydman (Pechmann & Frydman, 2013). The tAIg value for each gene was determined as the geometric mean of all codons, excluding the stop codon.

CSC values for budding yeast were taken from Carneiro et al. (Carneiro *et al.*, 2019) and only values derived from mRNA half-life measurements within the last 10 years were included (Becskei, Coller, Cramer, Gresham, Struhl, Weis). CSC values for human cells were taken from Wu et al. (Wu *et al*, 2019). Mad1 and Mad2 sequences from opisthokonts were taken from Vleugel et al. (Vleugel *et al*, 2012). The human Mad1 sequence was swapped for the canonical isoform (UniProt, Q9Y6D9, MD1L1_HUMAN), the *S. pombe* Mad1 sequence was shortened N-terminally by 13 amino acids to start with what is now considered the correct start codon (pombase.org). All sequences shorter than 600 amino acids were omitted for Mad1. Protein sequences were aligned using MAFFT (G-INS-i, using DASH and Mafft-homologs [100 homologs, E=1e-30]) (Katoh *et al*, 2019). Sequences from all species other than *S.c.*, *S.p.* and *H.s.* were deleted for the display of conservation shown in Fig. S8. The moving average of CSC values across 9 codons was plotted along the length of the aligned sequence. The null distribution of the moving average was obtained by randomizing the codon order 10,000 times. Observed values that deviated by more than 2 standard deviations from the null mean were marked with filled circles.

### Gene expression models

Protein noise predictions (Fig. 2C; S3B,C) were made by only considering stochastic mRNA and protein synthesis and degradation and ignoring cell growth and division. The coefficient of variation (CV = standard deviation / mean) for protein is calculated as:

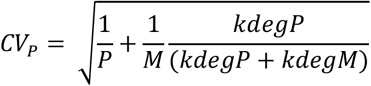

where P is the protein number per cell, M the mRNA number per cell, kdegM the mRNA degradation rate and kdegP the protein degradation rate (Swain, 2004). For the predictions in S3C, we assumed a protein number of 6,000 per cell, mRNA numbers of 1 to 1,000, and we varied RNA degradation rate in a range corresponding to half-lives of 1 to 60 minutes, and protein degradation rate in a range corresponding to half-lives of 15 to 600 minutes, which we consider a physiologically plausible range. Predictions were excluded when mRNA synthesis or protein synthesis rates became unrealistically high. We assumed this to be the case when mRNA synthesis rate was higher than 25 minute^−1^ or protein synthesis rate higher than 20 mRNA^−1^ minute^−1^. Assuming a gene with characteristics similar to a SAC gene (protein number = 6,000, mRNA number = 3.5, protein half-life = 360 minutes, mRNA half-life = 4 minutes) yields a CV prediction of 0.0575. In the figure, we labelled CV predictions less than 0.06 in light grey (low noise) and those equal or higher than 0.06 in dark grey (high noise).

The stochastic simulation of mRNA and protein numbers used the Gillespie algorithm in a Matlab script originally written by Daniel Charlebois and available on MathWorks (“Gillespie’s Direct Method Stochastic Simulation Algorithm“).

## Acknowledgments

We thank Tatiana Boluarte, Hunter Haynie, Jessica Malc, Haoyun Yang, Woong Sik Shin and Varun Gopala Krishna for their experimental help, Tony Carr’s lab for the yEGFP-containing plasmid, Yoshinori Watanabe’s lab for yeast strains, Daniel Zenklusen for the FISH protocol, as well as Andrea Ciliberto and all members of the Hauf Lab for comments on the manuscript. This work was supported by the National Science Foundation under grant no. 1616247.

## Author contributions

Conceptualization: EE, DW, SH; Formal analysis: EE, DW, JR, CM, SH; Funding acquisition: SH; Investigation: EE, DW, JR, CM, EKB, JC; Methodology: EKB, JC; Supervision: SH; Visualization: EE, DW, JR, SH; Writing – Original Draft Preparation: EE, DW, JR, SH; Writing – Review & Editing: EKB, JC.

## Conflict of Interest

None.

## Notes

### Competing Interest Statement

The authors have declared no competing interest.

