## Supplementary material for "Mitotic checkpoint gene expression is tuned by coding sequences": Figures S1-S8 and Tables S1-S5

**Figure S1**
**Esposito et al.**

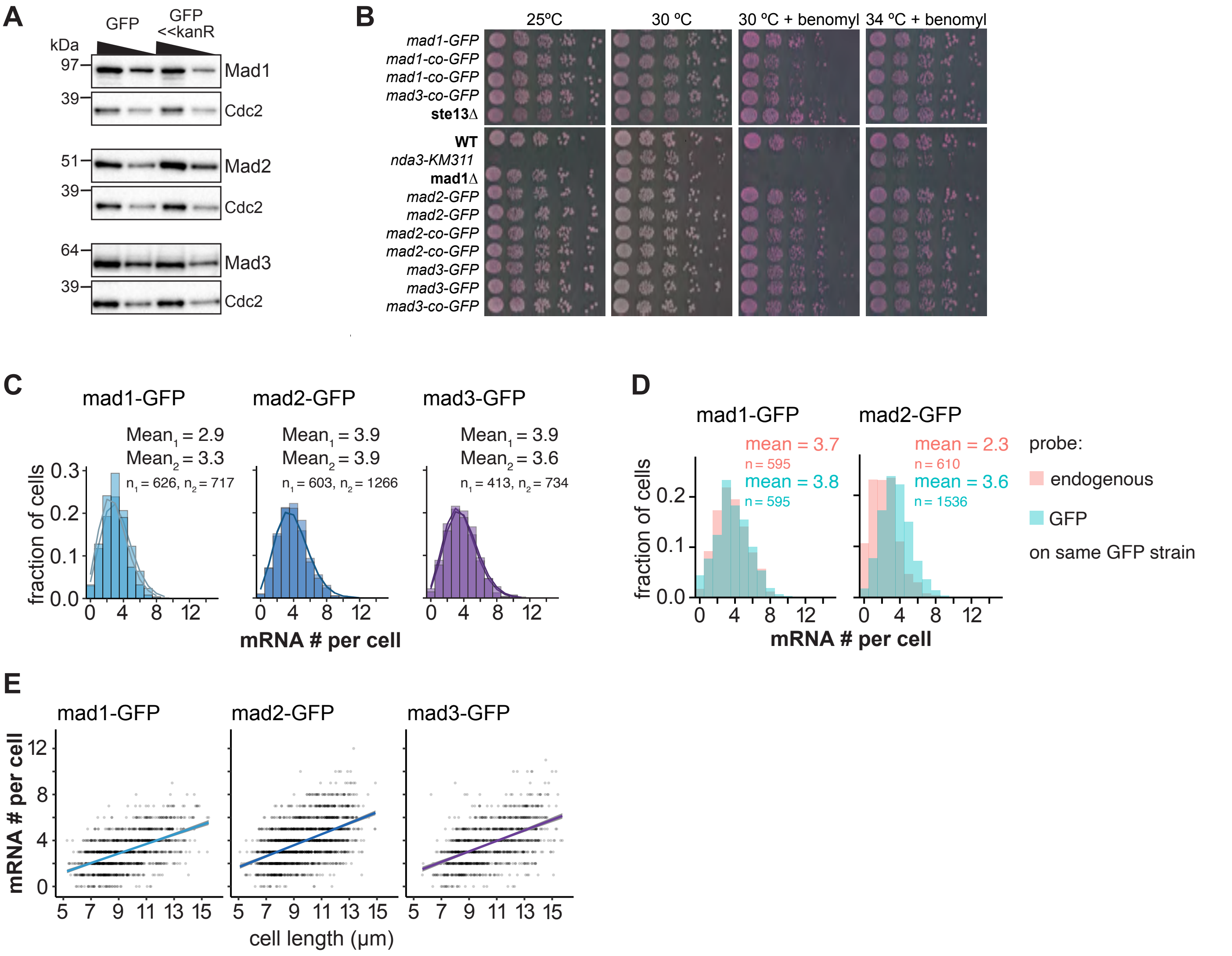


**Figure S1. Additional data on Mad1, Mad2, and Mad3 tagging and mRNA numbers.**
**(A)** Immunoblot comparing expression of *mad1*, *mad2* and *mad3* tagged at the endogenous locus either by marker-less insertion of yeast codon-optimized monomeric enhanced GFP (ymEGFP, here: GFP) or conventionally with GFP-S65T and a kanamycine-resistance cassette (GFP<<kanR). Antibodies against the endogenous proteins were used. Cdc2 was probed as loading control. A 1:1 dilution is loaded in the second lane for each sample.
**(B)** Growth assay for the indicated strains on rich medium plates without (left side) or with benomyl (right side).
**(C)** Frequency distribution of mRNA numbers per cell. Data from individual experiments which are shown combined in Figure 1. Probes were against the GFP portion of each fusion gene. Curves show fit to a Poisson distribution. The two fits for *mad2-GFP* overlap, so that only one line can be seen. **(D)** Frequency distribution of mRNA numbers per cell using probes against the endogenous gene or against GFP in strains expressing a GFP fusion of either *mad1* or *mad2*. The comparison illustrates that for Mad2-GFP either the endogenous probe is less sensitive, or there is considerable mRNA degradation from the 5' end leading to fewer detected spots with a probe on the endogenous gene than on the 3' end GFP tag. **(E)** Same experiments as in Figure 1 and (C). The mRNA number per cell is shown relative to the cell length. Lines show linear regression and grey area shows 95% confidence interval.

**Figure S2**
**Esposito et al.**

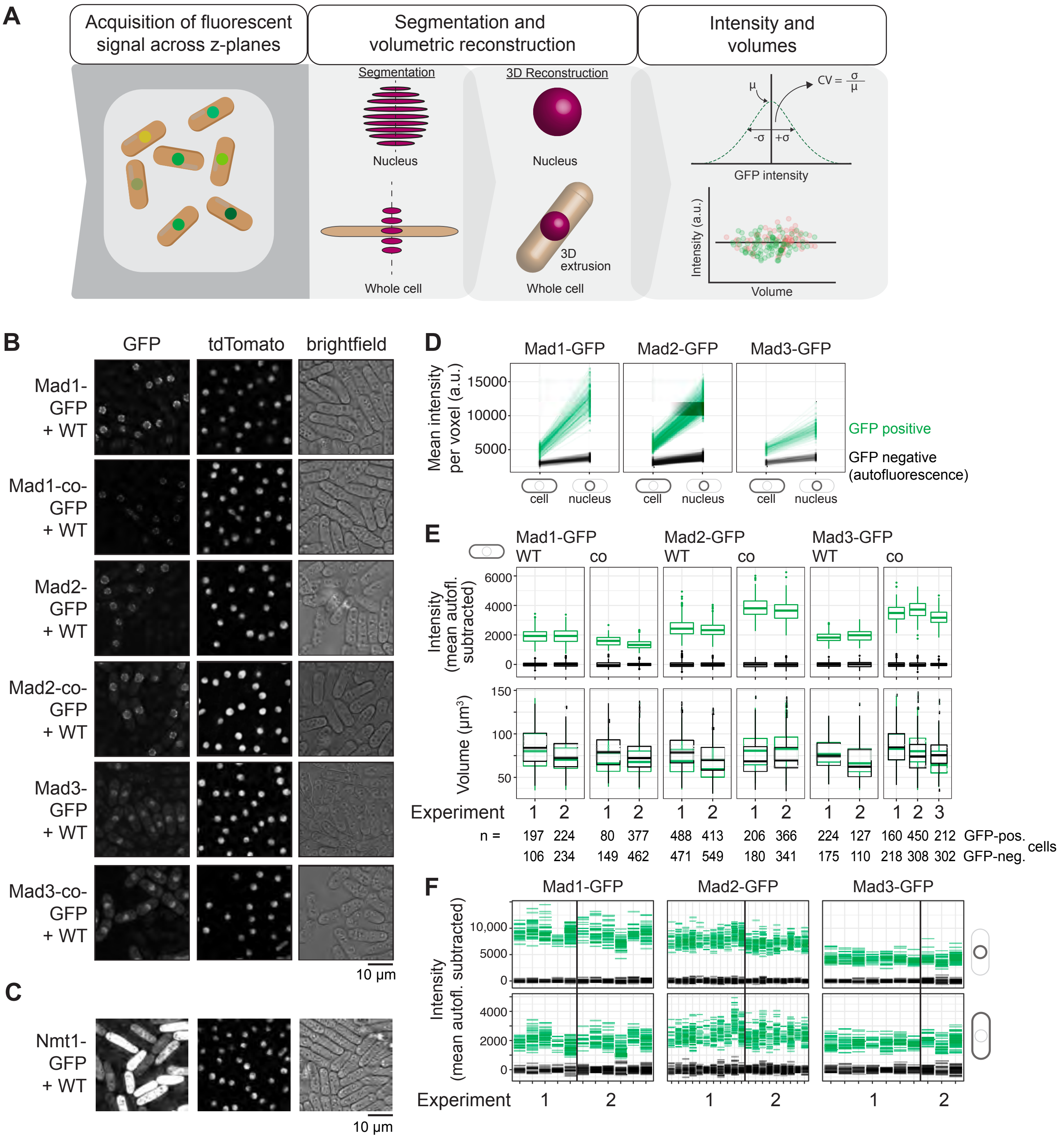


**Figure S2. Pomegranate pipeline image analysis to determine GFP intensities and noise.**
**(A)** Schematic overview of the Pomegranate pipeline (Baybay *et al.*, 2020): cells are segmented in two dimensions (2D) based on the brightfield image, nuclei are segmented in three dimensions (3D) based on tdTomato-NLS signal. To collect signal from the entire cell, the most in-focus 2D cell segmentation is extended into 3D by spherical extrusion. Signals are averaged from across all z-sections to obtain cellular and nuclear intensities. **(B, C)** Example pictures from live-cell imaging. The strains to be quantified are mixed with wild-type cells (WT, no GFP) to subtract autofluorescence. Cells are trapped in a microfluidics channel.
**(D)** GFP-expressing (green) and GFP-negative wild-type cells (black) are separated based on their distinct intensities in the green channel (a.u. = arbitrary units). For checkpoint proteins, k-means clustering with k = 2 was used. For Nmt1-GFP, the populations were split manually. **(E)** Volumes between GFP-positive (green) and GFP-negative (black) cells are similar, but intensities differ. For intensity, the mean signal intensity of the GFP-negative cells in each image was subtracted. (WT = wild-type coding sequence, co = codon-optimized coding sequence) **(F)** Fluorescence intensities of GFP-positive (green) and GFP-negative (black) cells in different images and experiments to demonstrate variability. Black vertical lines separate experiments on different days. Top: nuclear intensities; bottom: whole-cell intensities.

### Figure S3
### Esposito et al.

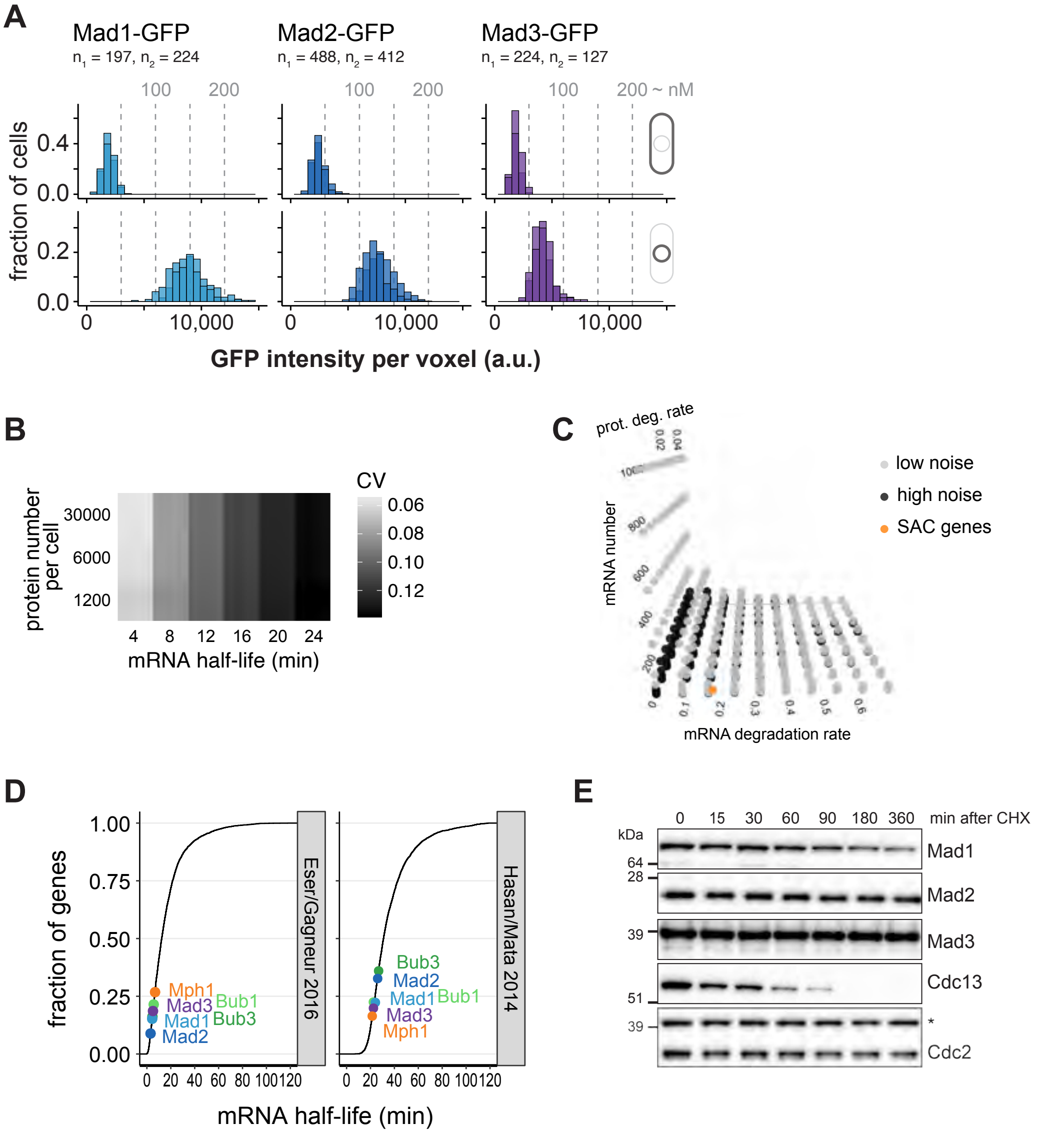


**Figure S3. Additional data on Mad1, Mad2, and Mad3 protein concentrations and half-life.**
**(A)** Concentration of GFP measured by live-cell imaging in whole cells (top) or nuclei (bottom). Two replicate experiments are overlaid. Estimates for concentration in nM are derived from comparison to previous absolute quantification of a mad3-GFP<<kanR strain and are very rough estimates only. **(B)** Protein noise (coefficient of variation; CV = std / mean) predictions from a simple gene expression model (see Methods for details). Protein half-life was set to 6 hours and protein synthesis rate was adjusted to reach each specified mean. Synthesis rate for mRNA was adjusted to maintain a mean mRNA number per cell of 3.5. **(C)** Simulated protein noise for different mRNA numbers, mRNA degradation rates and protein degradation rates. Noise was labeled as low when it was similar or lower than that of SAC genes and high otherwise. The mRNA degradation rate was varied in a range corresponding to half-lives of 1–60 minutes. The protein degradation rate was varied in a range corresponding to half-lives of 15–600 minutes. Missing points (e.g. high mRNA numbers at high mRNA degradation rates) are not included, because they would require non-physiologically high transcription or translation rates. The position where SAC genes are found in this grid is marked in orange. **(D)** Cumulative distribution of the mRNA half-lives of protein-coding *S. pombe* genes, measured by Eser *et al.* (2016) or Hasan *et al.* (2014). Position of spindle assembly checkpoint genes marked in color.
**(E)** Immunoblot of protein extracts harvested at the indicated times after translation shut-off by cycloheximide. One of the experiments quantified in Figure 2E.

**Figure S4**
**Esposito et al.**

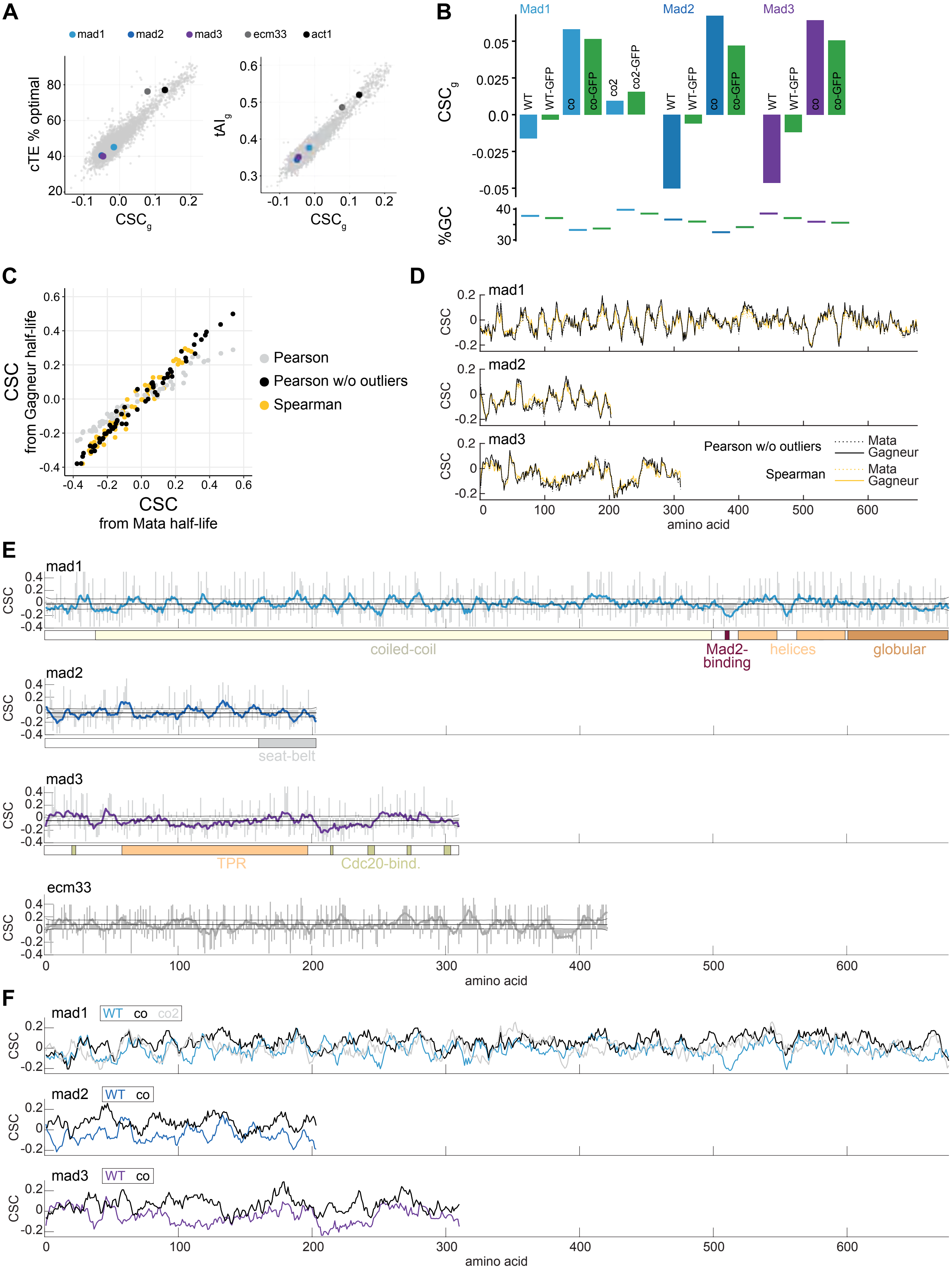

**Figure S4. Codon optimality of *mad1*, *mad2*, and *mad3* before and after codon-optimization.**
**(A)** Comparison between $CSC_g$ and other measures of codon optimality across protein-coding *S. pombe* genes. Genes relevant in this study are highlighted. **(B)** $CSC_g$ and GC content (%GC) for *mad1*, *mad2* and *mad3* before and after codon-optimization, and without or with taking the GFP tag into account. **(C)** CSC of the 61 amino acid-coding codons derived using mRNA half-life data from either the Mata group (Hasan *et al.*, 2014) or the Gagneur group (Eser *et al.*, 2016), and determined using different correlation methods (color), as explained in the methods section. **(D)** Moving average of the CSC across 9 codons along the length of each SAC gene, using codon CSC values derived from different datasets or with different correlation methods as in (E). **(E)** CSC value of each codon (grey bars) and moving average across 9 codons (colored line) along the length of each gene. For the SAC genes, functional and structural motifs are shown at the bottom. The middle black line represents the mean CSC across the gene (=$CSC_g$). The black lines above and below were obtained by randomly permuting the CSC values along the gene 10,000-times and determining the moving average across 9 codons (as for the original data). Shown is +/- 1 standard deviation of this randomized data. **(F)** Moving average of the CSC across 9 codons along the length of each SAC gene for both the wild-type and codon-optimized versions.

**Figure S5**
**Esposito et al.**

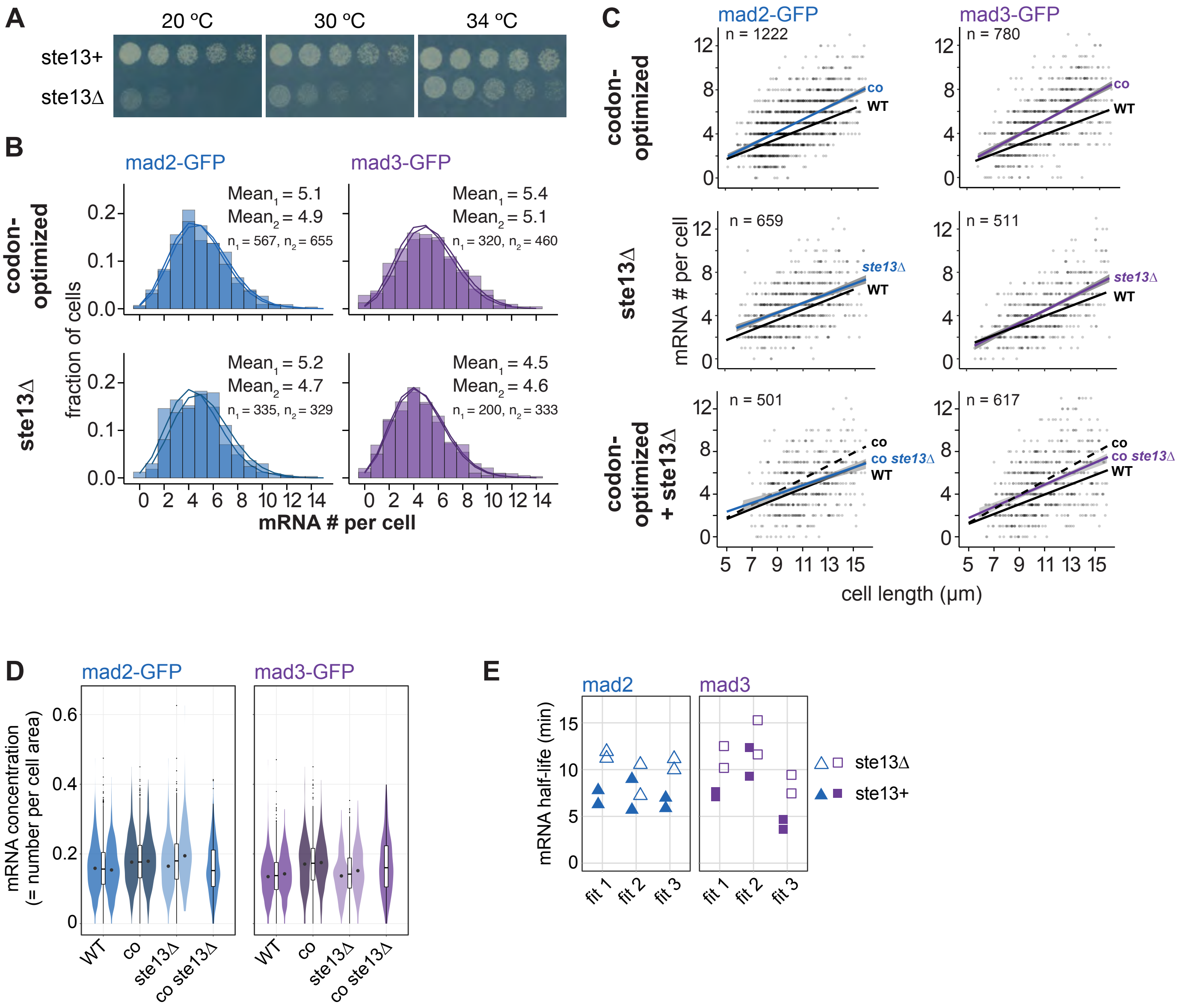


**Figure S5. Additional data on *mad2* and *mad3* mRNA numbers and half-lives after codon-optimization or *ste13*$^{+}$ deletion.**
**(A)** Growth assay for wild-type and *ste13*Δ cells on minimal medium plates. **(B)** Data from individual mRNA FISH experiments shown combined in Figure 3. Frequency distribution of mRNA numbers per cell. Probes were against the GFP portion of each respective fusion gene. Curves show fit to a Poisson distribution. **(C)** Same experiments as in Figure 3 and (B). The mRNA number per cell is shown relative to the cell length. Lines show linear regression and grey area shows 95% confidence interval. The black line shows the linear regression from wild type cells (WT, Figure S1). Only cells with lengths between 5 and 16 μm and widths between 2.2 and 5 μm were included. This excluded between 1 and 25 cells (or possibly falsely detected regions) in the different datasets. In the bottom panel, the linear regression from the codon-optimized gene data is additionally shown as dashed line. **(D)** Comparison of mRNA concentrations (calculated as mRNA number divided by cell area). Same data as in (B) and (C). Violin plots show data from different experiments with the median indicated (dot). Box plot shows the median and interquartile range from combining the different experiments. **(E)** Comparison of mRNA half-lives obtained using different fits to the same data (see Methods for details).

geneType

**Figure S6**
**Esposito et al.**

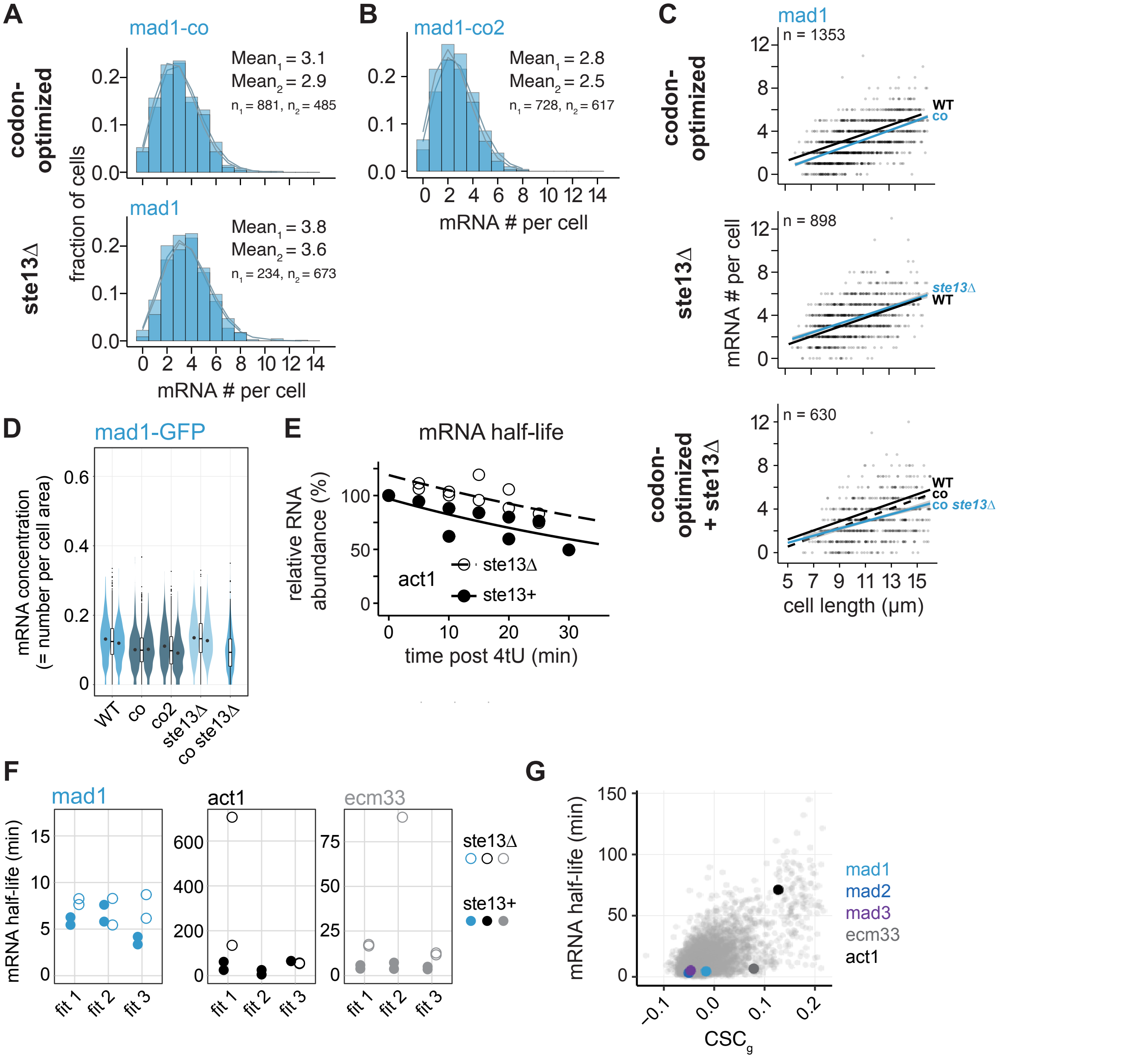


**Figure S6. Additional data on *mad1* mRNA number and half-life after codon-optimization or *ste13*⁺ deletion.**
**(A)** Data from individual mRNA FISH experiments shown combined in Figure 5. Frequency distribution of mRNA numbers per cell. Probes were against the GFP portion of *mad1-GFP*. Curves show fit to a Poisson distribution. **(B)** Frequency distribution of mRNA numbers per cell determined by mRNA FISH for a second codon-optimized version of *mad1*. Probes were against the GFP portion of *mad1-GFP*. Curves show fit to a Poisson distribution. **(C)** Same experiments as in Figure 5 and (A). The mRNA number per cell is shown relative to the cell length. Lines show linear regression and grey area shows 95% confidence interval. The black line shows the linear regression from wild type cells (WT, Figure S1). Only cells with lengths between 5 and 16 µm and widths between 2.2 and 5 µm were included. This excluded between 1 and 22 cells (or possibly falsely detected regions) in the different datasets. In the bottom panel, the linear regression from the codon-optimized gene data is additionally shown as dashed line. **(D)** Comparison of median mRNA concentrations (calculated as mRNA number divided by cell area). Same data as in (A) and (C). **(E)** Time course of mRNA abundances by qPCR following metabolic labeling and removal of the labeled pool (two independent experiments). Lines indicate fit (fit 3) to one-phase exponential decay, excluding the measurements at t = 0 in order to accommodate for non-instantaneous labeling by 4tU. The *ste13*⁺ data are the same as in Figure 2. **(F)** Comparison of mRNA half-lives obtained using different fits to the same data (see Methods for details). **(G)** $CSC_g$ values and mRNA half-lives (from Eser et al., 2016) for protein-coding *S. pombe* genes with the indicated genes highlighted.

### Figure S7
### Esposito et al.

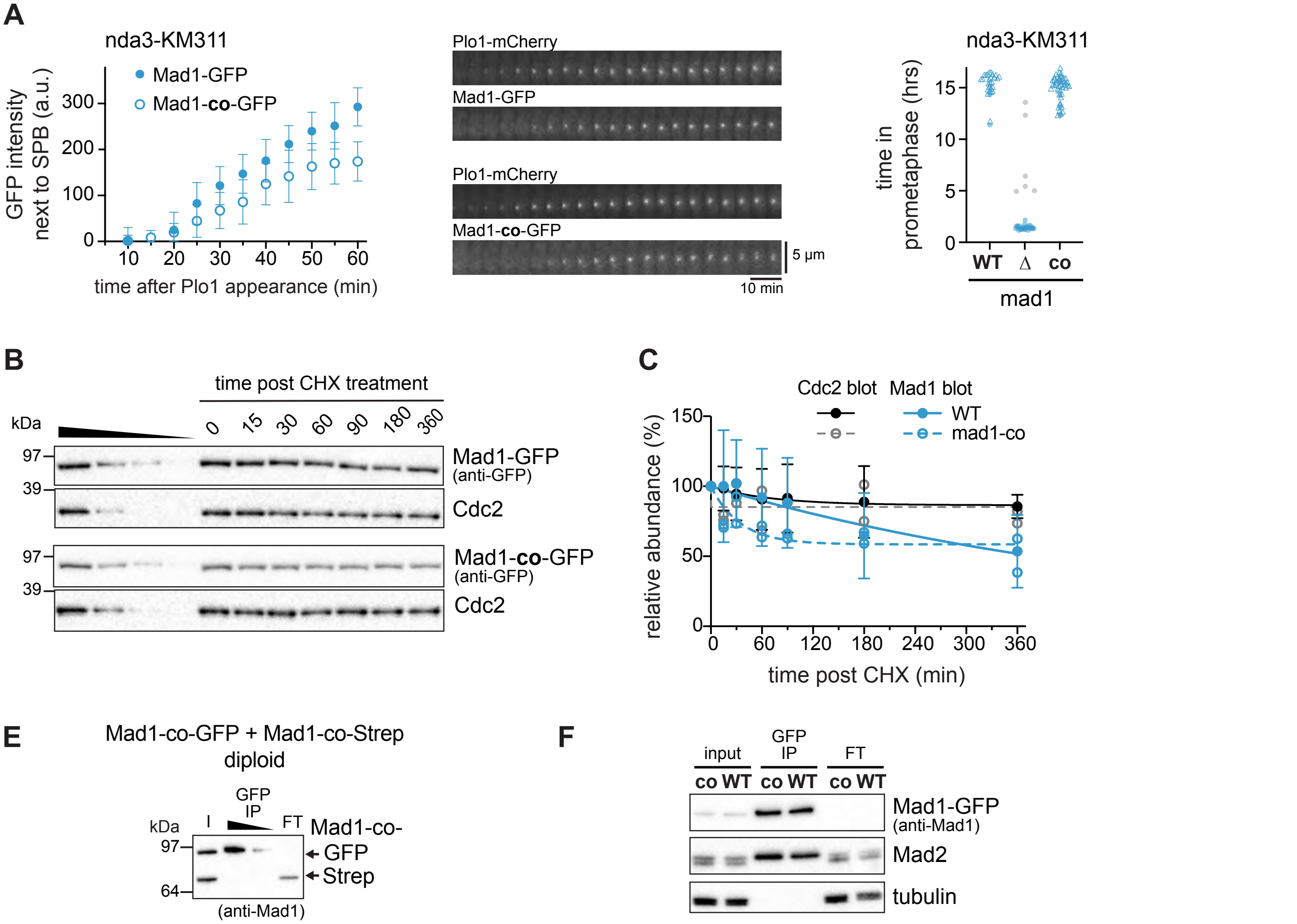


**Figure S7. Additional data on Mad1-co-GFP protein half-life and function.**
**(A)** Left and middle: The accumulation of Mad1-GFP at kinetochores in early mitosis was quantified in cells expressing the cold-sensitive tubulin mutant *nda3-KM311* and *plo1⁺-mCherry*, imaged at the restrictive temperature of 16 ºC. Mad1-GFP signals adjacent to spindle pole bodies (Plo1-mCherry) were interpreted as kinetochore localization (a.u. = arbitrary units; error bars = s.d.; n = 13 and 12 cells). One out of two experiments is shown. Right: Cells expressing the cold-sensitive tubulin mutant *nda3-KM311*, *plo1⁺-mCherry*, and the indicated versions of *mad1* were analyzed by live-cell imaging at the restrictive temperature of 16 ºC, similar to the experiment on the left. The time that each cell spent in prometaphase was determined by localization of Plo1 to spindle pole bodies (circle). Cells that had not yet exited mitosis when filming stopped are indicated by triangles and cells that died during mitosis by filled circles. n = 29, 32 and 53 cells. **(B)** Immunoblot of protein extracts at the indicated times after translation shut-off by cycloheximide. One of the experiments quantified in Figure 6F and (C). Mad1-GFP and Mad1-co-GFP were probed with anti-GFP. Cdc2 serves as control. **(C)** Same data as in Figure 6F, but now including one experiment for Mad1 WT cells, where levels for both Mad1 and Cdc2 at time points 15 min to 180 min were higher than at the 0 min time point. **(E)** Anti-GFP immunoprecipitation (IP) from extracts of diploid cells expressing Mad1-co-GFP and Mad1-co-Strep from the two endogenous loci. The membrane was probed with antibodies against Mad1. **I**, input; FT, flow-through. **(F)** Anti-GFP immunoprecipitation (IP) of Mad1-GFP (WT) and Mad1-co-GFP. Input, IP, and flow-through (FT) were probed with antibodies against Mad1, Mad2 and tubulin.

### Figure S8
### Esposito et al.

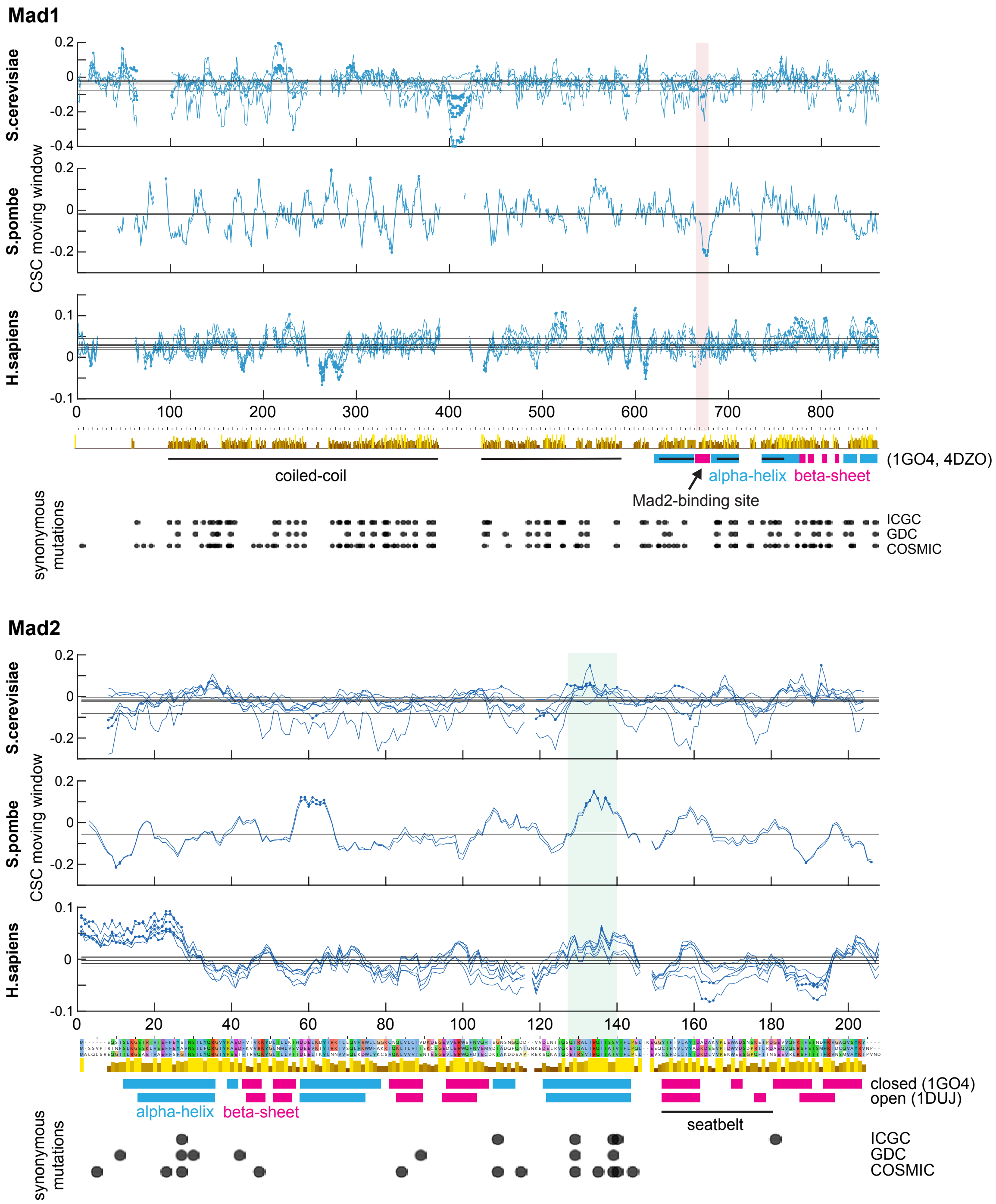


**Figure S8. CSC along Mad1 and Mad2 in *S. cerevisiae*, *S. pombe*, and *H. sapiens* and synonymous mutations observed in cancer.**
Mad1 and Mad2 were aligned using MAFFT. Yellow/brown bars beneath each panel indicate conservation. Blue lines show the moving CSC average across 9 codons. CSC values were derived from different mRNA half-life datasets, as explained in the Methods section, resulting in the multiple lines for each species. Filled circles indicate positions where the value deviates by more than two standard deviations from the mean for this particular dataset and gene. The thin horizontal lines indicate the mean of CSC values across the gene for each dataset. Secondary structure elements come from solved structures, whose protein data bank (PDB) identifiers are given in brackets. Red-shaded region: Mad2-binding site in Mad1 with markedly low CSC values in *S. pombe* and in one of the *S. cerevisiae* datasets. Green-shaded region: Conserved region in Mad2 with commonly high CSC values in these three organisms. Grey dots at the bottom indicate the position of synonymous mutations found in human cancer databases. In the ICGC and COSMIC data, synonymous mutations are enriched in the Mad2 region marked in green ($p < 0.005$ by chi-square test).

**Table S1 - Yeast strains**

| Strain Number | Mating type | Genotype |
|---|---|---|
| **Figure 1A, 2A; S2, S3A (protein concentrations and noise by live-cell imaging)** | | |
| SW184 | *h-* | *leu1 Z<<natR<<Padh31-tetR-tdTomato* |
| SW178 | *h+* | *mad1+-ymEGFP Z<<natR<<Padh31-tetR-tdTomato* |
| SW180 | *h+* | *mad2+-ymEGFP Z<<natR<<Padh31-tetR-tdTomato* |
| SW182 | *h+* | *mad3+-ymEGFP Z<<natR<<Padh31-tetR-tdTomato* |
| SW200 | *h?* | *nmt1+-ymEGFP<<kanMX6 Z<<natR<<Padh31-tetR-tdTomato* |
| **Figure 1B,C; S1C,E (mRNA numbers by smFISH)** | | |
| SW206 | *h+* | *mad1+-ymEGFP* |
| SW130 | *h+* | *mad2+-ymEGFP* |
| SW132 | *h+* | *mad3+-ymEGFP* |
| **Figure 1D (mRNA numbers by smFISH, tagged vs. untagged)** | | |
| SW642 | *h+* | *mad1+-ymEGFP* |
| SW130 | *h+* | *mad2+-ymEGFP* |
| JY001 | *h-* | wild type |
| **Figure 1F (test of mRNA co-localization)** | | |
| SW642 | *h+* | *mad1+-ymEGFP* |
| SW130 | *h+* | *mad2+-ymEGFP* |
| **Figure 2D (mRNA half-life)** | | |
| JY743 | *h-* | *leu1 ura4-D18* |
| ST932 | | *S. cerevisiae MET17pr-Fluc::lys2Δ (AMV54, Buchler lab)* |
| **Figure 2E; S3E (protein half-life)** | | |
| JY001 | *h-* | wild type |
| SU228 | *h+* | *bub1+-ymEGFP* |
| SL221 | *h-* | *leu1 ade6-M216 cut1+-GFP<<kanR* |
| **Figure 3D, S5B-D (mad2 and mad3 mRNA numbers by smFISH after codon-optimization and/or ste13 deletion)** | | |
| SW131 | *h+* | *mad2-codonOpt-ymEGFP* |
| SW133 | *h+* | *mad3-codonOpt-ymEGFP* |
| SW157 | *h+* | *mad2+-ymEGFP ste13Δ::kanMX6* |
| SW158 | *h+* | *mad3+-ymEGFP ste13Δ::kanMX6* |
| SW193 | *h+* | *mad2-codonOpt-ymEGFP ste13Δ::kanMX6* |
| SW194 | *h+* | *mad3-codonOpt-ymEGFP ste13Δ::kanMX6* |
| **Figure 3E, 5D; S5E (mad2 and mad3 mRNA half-life after ste13 deletion)** | | |
| JY743 | *h-* | *leu1 ura4-D18* |
| ST932 | | *S. cerevisiae MET17pr-Fluc::lys2Δ (AMV54, Buchler lab)* |
| SW239 | *h-* | *leu1 ura4-D18 ste13Δ::kanMX6* |
| **Figure 4A,C (Mad2 and Mad3 protein levels after codon-optimization by immunoblotting)** | | |
| SU257 | *h+* | *leu1+<< Park1-mCherry cut11+-mCherry<<hph mad2+-ymEGFP* |
| SU263 | *h+* | *leu1+<< Park1-mCherry cut11+-mCherry<<hph mad2-codonOpt-ymEGFP* |
| SU131 | *h-* | *leu1 ura4-D18 mad2+-ymEGFP* |
| SU128 | *h-* | *leu1 ura4-D18 mad2-codonOpt-ymEGFP* |
| SU294 | *h+* | *leu1+<< Park1-mCherry cut11+-mCherry<<hph mad3+-ymEGFP* |
| SW120 | *h+* | *leu1+<< Park1-mCherry cut11+-mCherry<<hph mad3-codonOpt-ymEGFP* |
| SW121 | *h+* | *leu1+<< Park1-mCherry cut11+-mCherry<<hph mad3-codonOpt-ymEGFP* |
| **Figure 4B,C (Mad2 and Mad3 protein levels after ste13 deletion by immunoblotting)** | | |
| JY001 | *h-* | wild type |
| JY002 | *h+* | wild type |
| SW148 | *h+* | *ste13Δ::kanMX6* |
| **Figure 4D,E; S2 (Mad2 and Mad3 protein levels after codon-optimization by live-cell imaging)** | | |
| SW184 | *h-* | *leu1 Z<<natR<<Padh31-tetR-tdTomato* |
| SW180 | *h+* | *mad2+-ymEGFP Z<<natR<<Padh31-tetR-tdTomato* |
| SW181 | *h+* | *mad2-codonOpt-ymEGFP Z<<natR<<Padh31-tetR-tdTomato* |
| SW182 | *h+* | *mad3+-ymEGFP Z<<natR<<Padh31-tetR-tdTomato* |
| SW183 | *h+* | *mad3-codonOpt-ymEGFP Z<<natR<<Padh31-tetR-tdTomato* |
| **Figure 5B; S6A-D (mad1 mRNA numbers by smFISH after codon-optimization and/or ste13 deletion)** | | |
| SW129 | *h+* | *mad1+-codonOpt-ymEGFP* |
| SW618 | *h+* | *mad1+-codonOpt2-ymEGFP* |
| SW156 | *h+* | *mad1+-ymEGFP ste13Δ::kanMX6* |
| SW192 | *h+* | *mad1-CodonOpt-ymEGFP ste13Δ::kanMX6* |
| **Figure 5C,D; S6E-F (mad1 and ecm33 mRNA half-life after ste13 deletion)** | | |

| | | |
|---|---|---|
| JY743 | *h-* | *leu1 ura4-D18* |
| ST932 | | *S. cerevisiae MET17pr-Fluc::lys2Δ (AMV54, Buchler lab)* |
| SW239 | *h-* | *leu1 ura4-D18 ste13Δ::kanMX6* |
| **Figure 6A,C (Mad1 protein levels after codon-optimization by immunoblotting)** | | |
| SW106 | *h+* | *leu1+<<Park1-mCherry cut11+-mCherry<<hph mad1+-ymEGFP* |
| SU282 | *h+* | *leu1+<<Park1-mCherry cut11+-mCherry<<hph mad1-codonOpt-ymEGFP* |
| SU499 | *h-* | *leu1 ura4-D18 mad1+-ymEGFP* |
| SU136 | *h-* | *ura4-D18 mad1+-codonOpt-ymEGFP* |
| **Figure 6B,C (Mad1 protein levels after ste13 deletion by immunoblotting)** | | |
| JY001 | *h-* | wild type |
| JY002 | *h+* | wild type |
| SW148 | *h+* | *ste13Δ::kanMX6* |
| **Figure 6D,E; S2 (Mad1 protein levels after codon-optimization by live-cell imaging)** | | |
| SW184 | *h-* | *leu1 Z<<natR<<Padh31-tetR-tdTomato* |
| SW178 | *h+* | *mad1+-ymEGFP Z<<natR<<Padh31-tetR-tdTomato* |
| SW179 | *h+* | *mad1-codonOpt-ymEGFP Z<<natR<<Padh31-tetR-tdTomato* |
| **Figure 6F; S7A,B (Mad1 protein half-life after codon-optimization)** | | |
| SW199 | *h?* | *mad1-codonOpt-ymEGFP mad2-codonOpt-ymEGFP mad3-codonOpt-ymEGFP* |
| SW234 | *h+* | *mad1+-ymEGFP mad2+-ymEGFP mad3+-ymEGFP* |
| **Figure 6G (Test for Mad1 co-translational assembly in haploid strain with tagged and untagged copies of mad1)** | | |
| SU164 | *h+* | *leu1<<110nt-mad1+-ymEGFP-164nt* |
| **Figure 6I (Test for Mad1 co-translational assembly in diploid strain with two differently tagged copies of mad1)** | | |
| SW623 | *h+/h-* | *leu1/leu1 ade6-M210/ade6-M216 mad1+-TEV-Strep2/mad1+-ymEGFP* |
| **Figure 7A,B (Replacement of mad1 coding sequence with GFP or GFP fusions)** | | |
| JY001 | *h-* | wild type |
| SW642 | *h+* | *mad1+-ymEGFP* |
| SW201 | *h-* | *mad1Δ::ymEGFP* |
| SW207 | *h+* | *mad1Δ::mad1(66bp)-ymEGFP* |
| SW221 | *h+* | *mad1Δ::mad1(108bp)-ymEGFP* |
| SU200 | *h?* | *mad1Δ::nmt1+-ymEGFP* |
| **Figure S1A (Immunoblot of strains traditionally tagged or tagged by scar-free genome editing)** | | |
| SW206 | *h+* | *mad1+-ymEGFP* |
| SK578 | *h+* | *leu1 mad1+-GFP<<kanR cut11+-mCherry<<hph* |
| SW130 | *h+* | *mad2+-ymEGFP* |
| SK580 | *h+* | *mad2+-GFP<<kanR cut11+-mCherry<<hph* |
| SW132 | *h+* | *mad3+-ymEGFP* |
| SK581 | *h+* | *mad3+-GFP<<kanR cut11+-mCherry<<hph* |
| **Figure S1B (Growth assay of strains with tagged mad1, mad2 and mad3, and ste13 deletion strain)** | | |
| SW206 | *h+* | *mad1+-ymEGFP* |
| SW129 | *h+* | *mad1-codonOpt-ymEGFP* |
| SW148 | *h+* | *ste13Δ::kanMX6* |
| JY001 | *h-* | wild type |
| PV273 | *h+* | *leu1 ade6-M216 nda3-KM311* |
| ST042' | *h-* | *leu1 ura4-D18 mad1Δ::ura4+* |
| SW130 | *h+* | *mad2+-ymEGFP* |
| SW131 | *h+* | *mad2-codonOpt-ymEGFP* |
| SW132 | *h+* | *mad3+-ymEGFP* |
| SW133 | *h+* | *mad3-codonOpt-ymEGFP* |
| **Figure S1D (Comparison of endogenous and GFP FISH probes)** | | |
| SW642 | *h+* | *mad1+-ymEGFP* |
| SW130 | *h+* | *mad2+-ymEGFP* |
| **Figure S5A (Growth assay of ste13 deletion strain)** | | |
| JY001 | *h-* | wild type |
| SW147 | *h-* | *ste13Δ::kanMX6* |
| **Figure S7A (Test for kinetochore association and checkpoint function of codon-optimized Mad1)** | | |
| SW601 | *h+* | *mad1-codonOpt-ymEGFP plo1+-mCherry<<natR nda3-KM311* |
| SW603 | *h-* | *mad1+-ymEGFP plo1+-mCherry<<natR nda3-KM311* |
| SW605 | *h-* | *mad1Δ::ymEGFP plo1+-mCherry<<natR nda3-KM311* |
| **Figure S7E (Test for Mad1 co-translational assembly in diploid strain with two differently tagged copies of mad1)** | | |
| SW666 | *h+/h-* | *leu1/leu1 ade6-M210/ade6-M216 mad1+-codonOpt-ymEGFP/mad1+-codonOpt-TEV-Strep2* |
| **Figure S7F (Test for Mad2-association of codon-optimized Mad1)** | | |
| SW129' | *h+* | *mad1-codonOpt-ymEGFP* |
| SW206 | *h+* | *mad1+-ymEGFP* |

**Table S2 - sgRNA targeting sequences**

| gene | targeting.sequence |
|---|---|
| mad1 | GCGCTCTCCGAGATTAGCAT |
| mad2 | ATTGGGTAGACAGTGACCCT |
| mad2 | GTTCAGGTGAAGATTTAGAG |
| mad3 | GCAATTTACTCACCGTTGGT |
| leu1 intergenic | GTAAGTACACAGCGACAACT |

| Table S3 - codon-optimized SAC gene sequences |
| --- |

| gene | company | sequence |
| --- | --- | --- |
| **mad1-codonOpt** | Genscript | ATGGCGGATTCTCCTAGaGATCCtTTCCAATCACGTTCACAATTACCTAGATTTTTAGCAACATCAGTTAAGAAAC<br>CAAACCTTAAGAAACCTTCTGTTAACTCTGCTAAtGGTATGTGAAACTTATATTTGACGTTTGTTTATGGATCTGAC<br>ACCCTGTAGAAACTAAAAATCCTAAGCTTGCTTCTCTTGAATTTCAATTGGAAAATCTTAAAAATGATCTTAAGCG<br>TAAGGAACTTGAATTTGAACGTGAACAAATTGAACTTCAACGTAAATTGGCTGAAGAACATGAACAAAAGAATTCT<br>TTGCAACTTCGTCTTACTTTGGTTGAAAAGCAACTTGAAGAACAATCTACTTCTTATCAAAAGGAAATTGAAGAAG<br>TTCGTAATGAAAAAGAAGCTACTCAAGTTAAGATTCATGAACTTCTTGATGCTAAGTGGAAGGAAATTGCTGAATT<br>GAAGACTCAAATTGAAAAGAATGATCAAGCTCTTTCTGAAAAGAATCATGAAGTTATGGTTTCTAACCAAGCTTTG<br>CAAATGAAGGATACTAATCTTACTAATTTGGAAAAGCTTTTTGCTGATTCTCGTGAACAATTGGAAACTAAGTGTA<br>AGGAATTAGCTGCTGCTGAACAACAACTTCAAGAATTGTCTGTTCATAATCAACAACTTGAAGAATCTATTAAACA<br>AGTTTCTTCTTCTATTGAATTGGAAAAGATTAATGCTGAACAACGTCTTCAAATTTCTGAACTTGAAAAGCTTAAA<br>GCTGCTCAAGAAGAACGTATTGAAAAGCTTTCTTCTAACAACCGTAATGTTGAAATTCTTAAAGAAGAAAAGAATG<br>ATCTTGAATCTAAGCTTTACCGTTTTGAAGAATACCGTGATAAGGTTGCTACTCTTGAACTTGAAAACGAAAAGAT<br>TCAAACTGAACTTAACTCTTGGAAATCTTTGATTACTAATGAATTACCTACTCCTGAAGCTGTTTCTAATAAGTTG<br>GTTTTTCTTCAAAACACTAATGCTAATCTTGGTGAACGTGTTTCTTCTTTGGAATCTCAACTTTCTAATAAGCCTGC<br>TAATCAACCTCTTGGTGCTAATGAAAAAGATGCTGCTCATATTACTGAATTGGAAACTAAGCTTAAGGAATTGCAT<br>GAACAAAATCGTCGTTTACAACGTCAAAAATCTCTTGCTACTCAAGAAATTGATTTGCTTCGTGAAAATTTGAAGT<br>CTTACGATGATGAAGAAGCTATTCTTTCTGAAAAGAATACTGATATGAAGAAACTTGAACGTATTGAAGGTTTAGT<br>TAAGCTTGTTGATGAATACAAGTTGAAATTAGAATCTATGCCTGTTTCTCTTGATGTTGATGAAACTTCTGATGAA<br>GTTTCTTTGCAAAAACGTCGTCGTAAAAATGAACATAAGGATGCTGGTTACGTTACTGAACTTTATCGTAAAAATC<br>AACATCTTTTGTTTCAAGTTAAAGAAAAGACTAACATTGAAGCTTTTCTTCGAGAACAAATTATTACTTTGGAATCT<br>TCTATTGCTACTCTTCGTCAAGAATTGGCTCAAGTTACTGAAATTAATTCTTGTCGTGTTCTTCAACATCGTTCTAA<br>CCCTACTCTTAAGTACGAACGTATTAAAGCTGCTCAATTGGAAATGCTTAACGCTGAAAACTCTGCTCTTAAGGC<br>TCTTCTTGAAGATAAGAAAGTTGATTGTCTTCCTATTCAATCTTTTAAAATTGCTGAACGTAAAGCTCTTGATCTTA<br>AGAAAGAAGTTGCTGAACGTGAAAAACGTATTCAACGTCTTAAGGAAATTTTCTCTGTTAAGTCTTTGGAATTTCG<br>TGAAGCTGTTTTCTCTCTTTTTGGTTACAAGTTGGATTTTATGCCTAACGGTTCTGTTCGTGTTACTTCTACTTACT<br>CTCGTGAAGATAACACTGCTTTTATTTTTGATGGTGAATCTTCTACTATGAAATTGGTTGGTAATCCTTCTGGTCC<br>TGAATTTGAACGTTTAATTCGTTTTTGGTGTGATGAACGTAAAACTATTCCTGGTATGCTTGCTGCTCTTACTCTT<br>GAACTTCTTGATAAAAATGAT |
| **mad1-codonOpt2** | IDT | ATGGCAGACTCTCCTCGCGACCCCTTTCAATCCAGATCACAATTACCCAGATTCCTTGCAACTAGTGTTAAGAAA<br>CCTAATCTTAAAAAACCCTCAGTTAATTCTGCTAATGGTATGTGAAACTTATATTTGACGTTTGTTTATGGATCTGA<br>CACCCTGTAGAAACAAAAAATCCTAAATTAGCATCACTTGAGTTTCAGTTGGAAAACTTAAAAAATGATTTGAAGC<br>GCAAGGAACTTGAGTTTGAACGTGAGCAGATTGAGTTGCAACGCAAATTGGCAGAAGAACACGAGCAAAAAAAT<br>TCATTGCAGTTAAGATTGACCTTAGTGGAAAAGCAATTGGAAGAGCAATCAACATCTTATCAAAAGGAGATCGAG<br>GAAGTGAGAAACGAGAAAGAAGCTACTCAAGTAAAGATTCACGAGTTACTTGATGCCAAATGGAAAGAAATTGCT<br>GAGTTAAAAACCCAAATAGAAAAGAATGATCAGGCACTTTCCGAAAAAAATCACGAGGTAATGGTATCCAATCAA<br>GCCTTGCAGATGAAGGATACGAATCTTACAAACTTAGAAAAGTTATTTGCTGATTCTCGCGAGCAGTTAGAGACC<br>AAATGTAAAGAGTTAGCTGCTGCCGAGCAACAATTACAGGAATTATCTGTACATAACCAACAATTGGAAGAGTCT<br>ATCAAACAGGTATCTTCTTCCATCGAATTGGAGAAAATTAATGCTGAGCAACGATTACAGATCTCAGAGTTAGAG<br>AAACTTAAAGCAGCACAAGAAGAGCGAATTGAGAAGCTTTCATCAAACAATCGCAACGTAGAGATTTTGAAGGA<br>GGAGAAAAACGATTTAGAGAGTAAGTTGTACCGATTTGAAGAGTATCGAGATAAGGTAGCTACTTTGGAATTGGA<br>GAACGAGAAAATCCAGACAGAGTTGAATAGTTGGAAAAGTCTTATAACTAACGAGTTGCCCACTCCCGAAGCTG<br>TCAGTAATAAGCTTGTATTCTTGCAGAATACCAACGCAAATTTAGGAGAGCGAGTATCCTCTTTAGAGTCACAGT<br>TGTCAAATAAGCCTGCCAACCAACCCTTGGGTGCCAATGAAAAGGACGCTGCCCACATCACCGAGTTGGAAACA<br>AAGCTTAAGGAATTACACGAGCAAAACAGACGTCTTCAACGACAAAAGAGTTTGGCTACTCAGGAGATTGACTT<br>GTTGAGAGAAAATTTAAAATCATATGATGATGAAGAGGCCATACTTTCAGAGAAAAATACCGATATGAAAAAATTG<br>GAACGCATCGAAGGCTTAGTAAAACTTGTAGATGAGTACAAATTGAAGCTTGAATCAATGCCCGTCTCTCTTGAC<br>GTGGATGAAACTAGTGACGAAGTGTCATTACAAAAGCGAAGACGTAAAAACGAACATAAAGACGCTGGCTATGT<br>GACGGAGTTGTACCGCAAAAACCAACACTTGTTGTTTCAGGTGAAAGAGAAGACAAATATAGAAGCCTTCTTACG<br>AGAACAAATCATTACGTTGGAATCCTCAATCGCTACATTAAGACAGGAGCTTGCTCAGGTAACGGAGATAAACAG<br>TTGTCGCGTCCTTCAGCATCGTTCCAATCCCACTTTGAAGTACGAAAGAATAAAGGCTGCACAACTTGAGATGTT<br>GAACGCAGAAAACAGTGCTCTTAAGGCCTTGTTGGAGGATAAGAAGGTGGACTGTCTTCCCATCCAGAGTTTTA<br>AGATAGCTGAAAGAAAAGCATTGGATTTGAAAAAGGAAGTTGCAGAGAGAGAAAAACGCATTCAGCGTTTAAAG<br>GAGATCTTTTCAGTCAAATCCTTAGAGTTCCGTGAGGCTGTCTTTTCATTATTTGGATATAAGTTGGACTTCATGC<br>CTAACGGAAGTGTGCGTGTAACTTCCACGTATTCTCGAGAGGACAATACCGCTTTTATATTCGATGGAGAGTCAT<br>CAACCATGAAGTTGGTGGGTAACCCATCTGGACCAGAATTCGAGCGATTGATACGATTCTGGTGTGATGAGAGA<br>AAAACCATTCCTGGCATGTTAGCCGCTTTGACCCTTGAGTTATTAGATAAAAATGAC |

**Table S3 - codon-optimized SAC gene sequences (cont.)**

| | | |
|---|---|---|
| **mad2-codonOpt** | Genscript | ATGTCTTCTGTTCCTATTCGTACTAACTTTTCTCTTAAGGGTTCTTCTAAACTTGTTTCTGAATTTTTTGGTTTGTAA<br>ATGTTTCAAATATGCCATGCTAACTTAATGAAGAATACGCTGTTAACTCTATTTTGTTTCAACGTGGTATCTATCCT<br>GCTGAAGATTTTAAGGTTGTTCGTAAGTACGGTTTAAACATGCTTGTTTCTGTTGATGAAGAAGTTAAAACTTATA<br>TTCGTAAAATTGTTTCTCAATTACACAGTGAGTTGTTGTTCTATTATCACTTTTGGAATTAACAAATTTTTAGAATG<br>GATGTTTGCTAAGAAAATTCAAAAGCTTATTCTTGTTATTACTTCTAAGTGTTCTGGTGAAGATTTGGAACGTTGG<br>CAATTCAATGTTGAAATGGTTGATACTGCTGATCAATTTCAAAACATTGGTAATAAGGAAGATGAACTTCGTGTTC<br>AAAAGGAAATTCAAGCTCTTATTCGTCAAATTACTGCTACTGTTACTTTTCTTCCTCAACTTGAAGAACAATGTACT<br>TTTAATGTTTTAGTTTATGCTGATAAAGATTCTGAAGTTCCTACTGATTGGGTTGATTCTGATCCTCGTATTCTTCG<br>TGATGCTGAACAAGTTCAATTGCGTTCTTTTTCTACTTCTATGCATAAGATTGATTGTCAAGTTGCTTATCGTGTTA<br>ATCCT |
| **mad3-codonOpt** | IDT | ATGGAACCACTTGATGCAGGTAAAAACTGGGTTCATATGGATGTTATTGAACAATCTAAGGAAAACATTGAACCA<br>CGAAAGGCAGGTCATTCTGCTTCTGCTTTGGCTAAGTCTTCTTCTCGTAACCATACTGAAAAAGAAGTTGCTGGT<br>TTACAAAAAGAACGTATGGGTCATGAGCGTAAGATTGAAACTTCTGAATCTCTTGATGATCCTTTGCAAGTTTGG<br>ATTGATTACATTAAGTGGACTCTTGATAACTTTCCTCAAGGTGAAACTAAGACTTCTGGTTTGGTTACTTTACTTG<br>AACGTTGTACTCGTGAATTTGTTCGTAACCCTTTGTACAAGGATGATGTTCGTTATCTTCGTATTTGGATGCAATA<br>CGTTAACTACATTGATGAACCTGTTGAATTATTTTCTTTTCTTGCTCATCATCATATTGGTCAAGAATCTTCTATTT<br>TCTATGAAGAATACGCTAACTACTTTGAATCTCGTGGTTTATTTCAAAAGGCTGATGAAGTTTACCAAAAGGGTAA<br>ACGTATGAAGGCTAAGCCTTTTCTTCGTTTTCAACAAAAGTACCAACAATTCACTCATCGTTGGCTTGAATTTGCT<br>CCTCAATCTTTTTCTTCTAACACTAACTCTGTTAACCCTCTTCAAACTACTTTTGAATCTACTAACATTCAAGAAAT<br>TTCTCAATCTCGTACTAAGATTTCTAAGCCTAAGTTTAAATTTTCTGTTTATTCTGATGCTGATGGTTCTGGTAAAG<br>ATGGTCAACCTGGTACTTGGCAAACTTTGGGTACTGTTGATCAACGTCGTAAGGAAAACAACATTTCTGCTACTT<br>CTTGGGTTGGTGAAAAATTGCCTTTAAAATCTCCTCGTAAGTTAGATCCATTGGGAAAGTTTCAAGTTCATTGCG<br>ATGAAGAAGTTTCAAAGGAA |

**Table S4 - FISH probes**

| mad2 | ymEGFP | mad1 |
|---|---|---|
| cgttcttatgggaacgctag | cagtgaataattcttcacctt | gaacggatccctaggagaatc |
| gtttggaagaacctttcagt | tcaaccaaaattgggacaaca | aatctaggcaactgtgaacgc |
| aaaggattgagttcaccgca | gaccattaacatcaccatcta | atttggtttcttaacgcttgt |
| gtcttctgctgggtaaattc | ccttcaccggagacagaaaat | gcagaattaacagaaggcttc |
| tccatatttccgaacaactt | gtcaatttaccgtaagtagca | agctagtttgggatttttgt |
| tcatctacactgacaagcat | atggaactggcaatttaccag | ccttccgctttaaatcatttt |
| tcgaatgtaagtcttgacct | aaagtagtgactaaggttggc | actcaatttgttcacgctcaa |
| ttgcaaacatccatttgtgt | tgttgtttcatatgatctggg | ttcttctgcaagtttctttg |
| cgctctaaatcttcacctga | ctggcatggcagacttgaaaa | gctgtaacgaattcttctgtt |
| catctccacattaaactgcc | gttctttcttgaacataacct | agttgcttttcaactagagtt |
| gaaattgatcagctgtgtca | agttaccgtcatctttgaaaa | gataagaagtagactgctcct |
| atcttctttgttgccaatgt | ttgacttcagctctggtcttg | ccttttcatttcttacttctt |
| tttctttttgtactcgcagt | taactaaggtatcaccttcaa | ttgcatcaagtagttcatgga |
| tgtagcagtgatttgacgaa | ataccttttaattcgattcta | ttcaactctgcaatctctttc |
| ctagttgaggcaaaaaggtc | cctaaaatgttaccatcttct | gatcattcttttctatctggg |
| acgttaaacgtacactgttc | gagagttatagttgtattcca | acttcatgattcttttcactt |
| gtctttatcagcgtatacca | gtcagccatgatgtaaacatt | tgcaaagcttgatttgagacc |
| cccaatctgttggaacttcg | actttgataccattcttttgt | gtttgtaagattggtatcctt |
| aaaatcctagggtcactgtc | gttgtgtctaattttgaagtt | ggaatccgcaaagagttttc |
| ttgaacttgttcagcatctc | attgaacagaaccatcttcaa | ttccttacacttcgtttcaag |
| atttgtgcatactcgtact | ttttgttgataatggtcagct | gaaagctcttgtaattgctgc |
| ggattcactcgatatgcaac | agactggaccatcaccaattg | tcttccaattgctgattatga |
| | aagtaatggttgtctggtaac | aactagaaacctgcttgatgg |
| | ggataaacgagattgagtgga | gcatttatttttccagttca |
| | ctctcttttcgtttggatctt | cgctaatttgaagacgttgct |
| | aattctaacaagaccatgtgg | cgcagcttttaattttccaa |
| | ggtaataccagcagcagtaac | gatagcttttcaattcgttct |
| | ttgtacaattcatccatacca | ccttgagaatttcaacattcc |
| | | ttggactccaaatcgtttttc |
| | | aaccttatccctatattcttc |
| | | tcattttcgagttcaagggta |
| | | gggttggtaactcgttagtaa |
| | | acgagtttgtttgaaacagct |
| | | ttagcattggtattctgtaga |
| | | taaactagaaacgcgctctcc |

**Table S5 - qPCR primers**

| Target | Forward | Reverse | |
|---|---|---|---|
| act1 | CCAAATCCAACCGTGAGAAGA | GTACGACCAGAGGCATACAAAG | |
| cdc2 | GGTATCGTGCTCCTGAAGTATTG | CAGAGTCACCGGGAAATAATGG | |
| ecm33 | ATCATTCGCTCTCACTCTTCTTT | GTACCTTGGGCGGAGATATTG | |
| mad1 | CCTAATGGGAGTGTTCGTGTTA | CCTGATGGATTACCAACCAATTTC | |
| mad2 | TTAGAGCGGTGGCAGTTTAAT | CTCGCAGTTCATCTTCTTTGTTG | |
| mad3 | CGGATGGTTCTGGAAAGGAT | CTACCCAAGAAGTAGCCGATATG | |
| S.c. ACT1 | TGGATTCCGGTGATGGTGTT | TCAAAATGGCGTGAGGTAGAGA | Chan et al. 2018 |
| ymEGFP | TGAAGGTGAAGGTGATGCTAC | CTAAGGTTGGCCATGGAACT | |
